# A genome-wide innateness gradient defines the functional state of human innate T cells

**DOI:** 10.1101/280370

**Authors:** M. Gutierrez-Arcelus, N. Teslovich, A. R. Mola, H. Kim, S. Hannes, K. Slowikowski, G. F. M. Watts, M. Brenner, S. Raychaudhuri, P. J. Brennan

## Abstract

Innate T cells (ITCs), including invariant natural killer T (iNKT) cells, mucosal-associated invariant T (MAIT) cells, and γδ T cell populations, use conserved antigen receptors generated by somatic recombination to respond to non-peptide antigens in an innate-like manner. Understanding where these cells fit in the scheme of immunity has been a puzzle since their discovery. Here, immunophenotyping of 101 individuals revealed that these populations account for as much as 25% of peripheral human T cells. To better understand these cells, we generated detailed gene expression profiles using low-input RNA-seq and confirmed key findings through protein-level and functional validation. Unbiased transcriptomic analyses revealed a continuous ‘innateness gradient’ with adaptive T cells at one end followed by MAIT, iNKT, Vδ1^+^ T, Vδ2^+^ T, and natural killer cells at the other end. Innateness was characterized by decreased expression of translational machinery genes and reduced proliferative potential, which allowed for prioritization of effector functions, including rapid cytokine and chemokine production, and cytotoxicity. Thus, global transcriptional programs uncovered rapid proliferation and rapid effector functions as competing goals that respectively define adaptive and innate states.

**One Sentence Summary:** Adaptive and innate T cells align along a continuous innateness gradient, reflecting a trade-off between effector function and proliferative capacity.

## Introduction

Within the spectrum of immune defense, “innate” and “adaptive” refer to pre-existing and learned responses, respectively. Mechanistically, innate immunity is largely ascribed to ‘hardwired,’ germline-encoded immune responses, while adaptive immunity derives from recombination and mutation of germline DNA to generate specific receptors that recognize pathogen-derived molecules, such as occurs in T and B cell receptors. However, the paradigm that somatic recombination leads only to adaptive immunity is incorrect.

Over the past 15 years, T cell populations have been identified with T cell antigen receptors (TCRs) that are conserved between individuals. Many of these effector-capable T cell populations are established in the absence of pathogen encounter. Examples of such T cell populations include invariant natural killer T (iNKT) cells, mucosal-associated invariant T (MAIT) cells, γδ T cells, and other populations for which we have a more limited understanding (*1-3*). These “donor unrestricted” T cell populations have been estimated to account for as much as 10-20% of human T cells (*4*), and have critical roles in host defense and other immune processes. The existence of innate-like T cells suggests that somatic recombination, the machinery of adaptive immunity, is working to generate TCRs that function as innate antigen receptors. We and others now refer to these cells as innate T cells (ITCs).

Most ITC populations share several important features. First, they do not recognize peptides presented by MHC class I and class II. iNKT cells recognize lipids presented by a non-MHC encoded molecule named CD1d (*5-7*). MAIT cells recognize small molecules including bacterial vitamin B-like metabolites presented by another non-MHC encoded molecule, MR1 (*8, 9*). It is not known whether specific antigen presenting elements drive the development of γδ T cells. One major γδ T cell population bearing V*γ*2-Vδ9 TCRs (Vδ2) is activated by self and foreign phospho-antigens in conjunction with a transmembrane butyrophyllin-family receptor, BTN3A1 (*10-13*). The antigens recognized by other human γδ T cell populations are not clear, although a subset of these cells recognizes lipids presented by CD1 family proteins (*14*), and recent data suggests that some may recognize butyrophyllin-family receptors (*15*). A second shared feature of ITCs is that their responses during inflammation and infection exhibit innate characteristics, such as rapid activation kinetics without prior pathogen exposure, and the capacity for antigen receptor-independent activation. Data in mice demonstrate that during diverse immune responses, including in host defense, cancer, autoimmunity, and allergic disease, a large portion of the iNKT cell pool is rapidly activated and orchestrates ensuing immune responses (*1, 16*). On the other hand, only a low frequency of the adaptive T cell pool responds during any given infection. Inflammatory cytokines such as IL-12, IL-18, and type I interferons can activate ITCs even in the absence of concordant signaling through their TCRs, and such TCR-independent responses have been reported in iNKT cells (*17, 18*), MAIT cells (*19, 20*), and γδ T cells (*21-24*). These ‘cytokine-only’ responses may explain how these cell populations contribute to immunity in diverse inflammatory contexts.

Given the similar functions reported among different ITC populations, we hypothesized that their effector capabilities might be driven by shared transcriptional programs. Here, we set out to transcriptionally define the basis of innateness in human ITCs by studying them as a group, focusing on their common features rather than what defined each population individually. We performed RNA-seq on highly-purified lymphocyte populations from the peripheral blood of six healthy donors in duplicate to generate robust transcriptional data, and confirmed key findings with protein-level and functional validation. Using unbiased methods to determine global inter-population relationships, we defined an ‘innateness gradient’ with adaptive cells on one end and natural killer (NK) cells on the other, in which ITC populations clustered between the prototypical adaptive and innate cells. Within the ITC cluster, populations segregated from adaptive to innate as MAIT, iNKT, Vδ1, and Vδ2. These data suggest that ITCs, with innate-like functionality and antigen receptors produced by the machinery of adaptive immunity, are a distinct family defined both transcriptionally and functionally. Interestingly, we observed decreased transcription of cellular translational machinery and a decreased capacity for proliferation as hallmarks of innate cells. Innate cells rather prioritized effector functions, including cytokine production, chemokine production, cytotoxicity, and reactive oxygen metabolism. Thus, growth potential and rapid effector function are hallmarks of adaptive and innate cells, respectively.

## Results

### Human immunophenotyping reveals high aggregate ITC frequencies

To characterize the abundance and variability of ITCs in humans, we quantified 4 major populations of innate T cells from 101 healthy individuals aged 20 to 58 years by flow cytometry, directly from peripheral blood mononuclear cells (PBMCs) in the resting state. We assessed the frequencies of iNKT cells, MAIT cells, and the two most abundant peripheral γδ T cell groups, those expressing a Vδ2 TCR chain (Vδ2) and those expressing a Vδ1 TCR chain (Vδ1). MAIT cells contributed from 0.1 to 15% of T cells (mean 2.4%), iNKT cells from undetectable to 1.1% (mean 0.09%), Vδ1 cells 0.25 −6.2% (mean 1.25%), and Vδ2 from 0.08 −22% (mean 4.7%). The sum of these 4 cell types accounted for 0.9 – 25.7% of an individual subject’s T cells (mean 8.4%) (Fig. 1A, Supplementary Table 1). Vδ2 cells were more abundant than Vδ1 in 82% of subjects, with the ratio of these two cell types ranging from 0.2 to 67.8 (mean 8.5). Age negatively associated with the total percentage of ITCs (P = 1.4e-05). MAIT (*r* = −0.42, P = 9.9e-06) and Vδ2 (*r* = −0.43, P = 4.7e-06) populations drove this association (Fig. S1A,B), even after accounting for the abundances of other cell types (P = 5.9e-04, P *=* 1.2e-04, respectively), which is consistent with previous findings (*25, 26*). We observed covariance between the frequencies of MAIT/iNKT cells (P = 0.02), corrected for the other cell types and age (Fig. S1C,D). We observed no significant associations between ITC percentage and gender, body mass index, or smoking status after accounting for age. Together, these results show human ITCs contribute a substantial portion of the peripheral T cell repertoire, are variable between individuals, and decrease with age.

**Figure 1.**
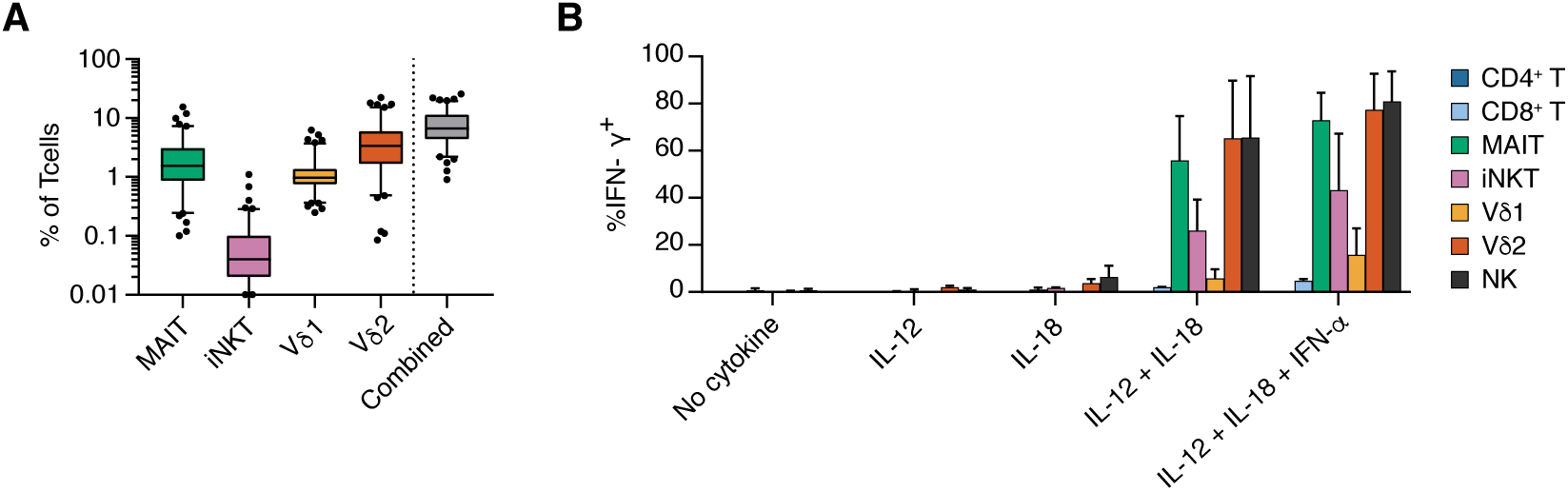
ITC immunophenotyping. (A) ITCs were quantified in 101 healthy donors by flow cytometry. The “Combined” group represents the sum of iNKT, MAIT, Vδ1, and Vδ2 T cells. For boxplots, 5-95 percentile and outliers are shown. (B) Intra-cellular staining for IFN-*γ* production following cytokine stimulation without TCR activation, N=4 independent donors, s.e.m.

### ITC populations rapidly release cytokines

We next tested innate T cell populations for two functional hallmarks of innate effectors, rapid cytokine production and TCR-independent activation. To assess rapid cytokine production potential, we activated healthy donor PBMCs with phorbol 12-myristate 13-acetate (PMA) and ionomycin for 4 hours, followed by intracellular staining for interferon-*γ* (IFN-*γ*) production. Between 35 and 85% of MAIT, iNKT, Vδ1, and Vδ2 T cells produced IFN-*γ* under these conditions, while a smaller percentage of adaptive CD4^+^ T and CD8^+^ T cells produced this cytokine (Fig. S2A,B). To test the relative capacity of these cell types to respond to inflammatory cytokines alone, we activated PBMCs with IL-12 + IL-18 or IL-12 + IL-18 + IFN-*α* for 16 hours, and assessed IFN-*γ* production during the final 4 hours of stimulation. 20-80% of iNKT, MAIT, Vδ2, and NK cells produced cytokines under these conditions, while only a tiny portion of adaptive cells responded (Fig. 1B, Fig. S2C,D). Taken together, these studies show that ITC populations rapidly produce cytokines, and can do so in response to inflammatory cytokines even in the absence of TCR signals. Notably, we observed the latter activation mechanism almost exclusively in ITC populations.

### RNA-seq profiling of ITCs reveals a continuous innateness gradient

To better understand the biological properties of human ITCs on a genome-wide scale, we profiled their transcriptomes with RNA-seq. Ultra-low input RNA-seq profiling using 1,000 cells per sample enabled high-depth sequencing of even relatively rare human lymphocyte populations. From 6 healthy individuals, we sorted in duplicate four subsets of ITCs: iNKT, MAIT (defined as MR1-5-OP-RU tetramer^+^), Vδ1 and Vδ2 cells (Supplementary Table 2). From the same individuals, we also sorted CD4^+^ and CD8^+^ T cells as comparator adaptive T cells and NK cells as comparator innate cells (Fig. S3). Using SmartSeq2 to create poly(A)-based libraries, we generated 25 base pair, paired-end libraries sequenced at a depth of 4-12 million read pairs (Fig. S4). After sequence mapping, we calculated tpm (transcripts per million) values for each gene. We considered 19,931 genes as expressed (tpm>3 in ≥10 samples), including 12,730 protein-coding, 183 T cell receptor genes, 3,261 long noncoding RNA (lncRNA), and other lowly-expressed genes (e.g. pseudogenes, Fig. S4C).

Principal component analysis identified the major axes of variation in gene expression (Fig. 2A). The first principal component separated the subsets by a continuous ‘innateness gradient’ with CD4^+^ and CD8^+^ T cells on one end, and NK cells on the other end (Fig. 2B). Ordered from adaptive to innate along the first principal component, MAIT, NKT, Vδ1 and Vδ2 clustered in between the adaptive cells and NK cells. We then identified genes associated with the rank order of each lymphocyte population in the innateness gradient (CD4^+^ T = 1, CD8^+^ T= 2, MAIT = 3, iNKT = 4, Vδ1 = 5, Vδ2 = 6, NK = 7), using linear mixed models. This analysis revealed 1,884 genes significantly associated with the innateness gradient (P < 2.5e-06=0.05/19,931, correcting for 19,931 tests), including protein coding and lncRNA genes (Fig. 2C, Supplementary Table 3). Hereafter we refer to positive and negative associations with the ranked gradient as associations with ‘innateness’ and ‘adaptiveness,’ respectively. We defined an ‘innateness score’ as the magnitude of the change in expression level by an increase of one in the gradient (the *β* of the gradient variable within our linear mixed model).

**Figure 2.**
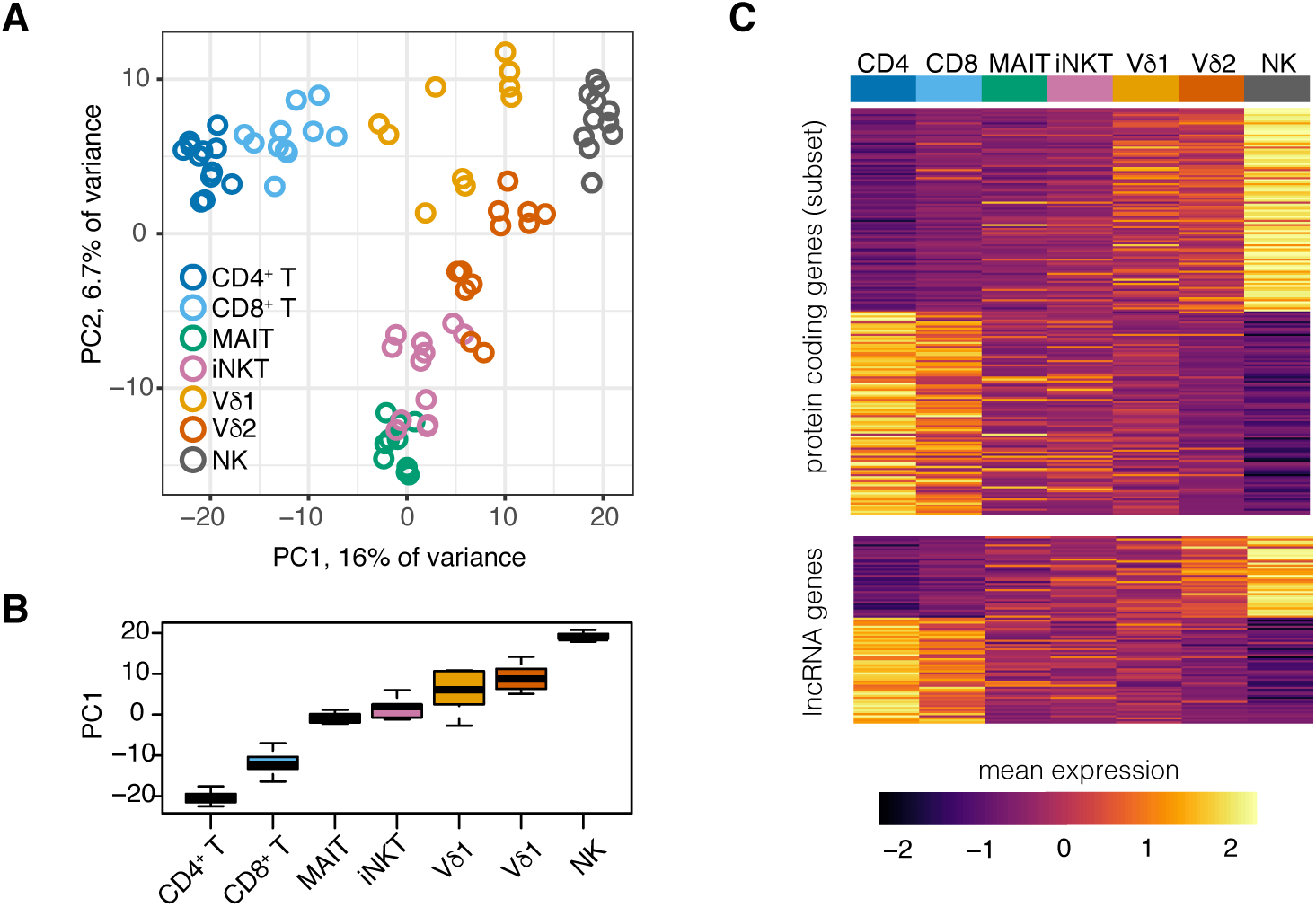
Transcriptomic profiling of ITCs reveals a continuous innateness gradient. (A) Principal component analysis performed on the top 1,022 most variable (s.d. > 1.4) and expressed genes. Plotted are scores for PC1 and PC2. (B) Distribution of PC1 scores for each cell type, arranged by rank order. (C) Heatplot of mean expression per cell type for genes associated with innateness gradient. Upper panel shows top 100 positive and negative significant associations within protein coding genes. Lower panel shows significant associations for 92 lncRNA genes. Genes within each heatplot were sorted by *β*. Gene expression level was scaled by row. N=6 donors, 1-2 replicates per cell type. Boxplots are described in methods.

### Associations with innateness: migration, cytotoxicity, cytokine production, and ROS metabolism

#### Cytotoxicity and chemokines

The Gene Ontology (GO) terms most associated with innateness included NK cell and lymphocyte chemotaxis, NK cell mediated immunity, cellular defense response, and several additional terms related to leukocyte migration and activation (Fig. 3A-D, specific GO terms indicated in figure legend and Fig. S5A). Using flow cytometry, we validated the expression of key genes, including killer cell lectin-like receptor (KLR) family genes and killer cell immunoglobulin-like receptor (KIR) genes (Fig. S5B). Cytotoxicity proteins such as perforin, granzyme B and granulysin also associated with innateness (Fig. 3E,F, Fig. S5B). Eight chemokines strongly associated with innateness, including *CCL3, CCL4, CCL5, XCL1* and *XCL2* (*P* < 9e-12), consistent with a role for innate lymphocytes in recruiting other inflammatory cell types to initiate inflammation. *IFNG* (the gene coding for IFN-*γ*) showed a significant association with innateness (P = 1.7e-06, Fig. 3G), and the baseline *IFNG* levels in each cell population predicted their production of IFN-*γ* upon stimulation (Fig. S2A,B). Since ITCs produce diverse cytokines and chemokines (*1, 2*), we quantified the total cytokine and chemokine transcriptome ‘mass’ in each cell type at baseline. We observed that the aggregate sum of the expression levels of the 37 cytokines and chemokine genes expressed in our dataset followed the innateness gradient (Fig. 3H).

**Figure 3.**
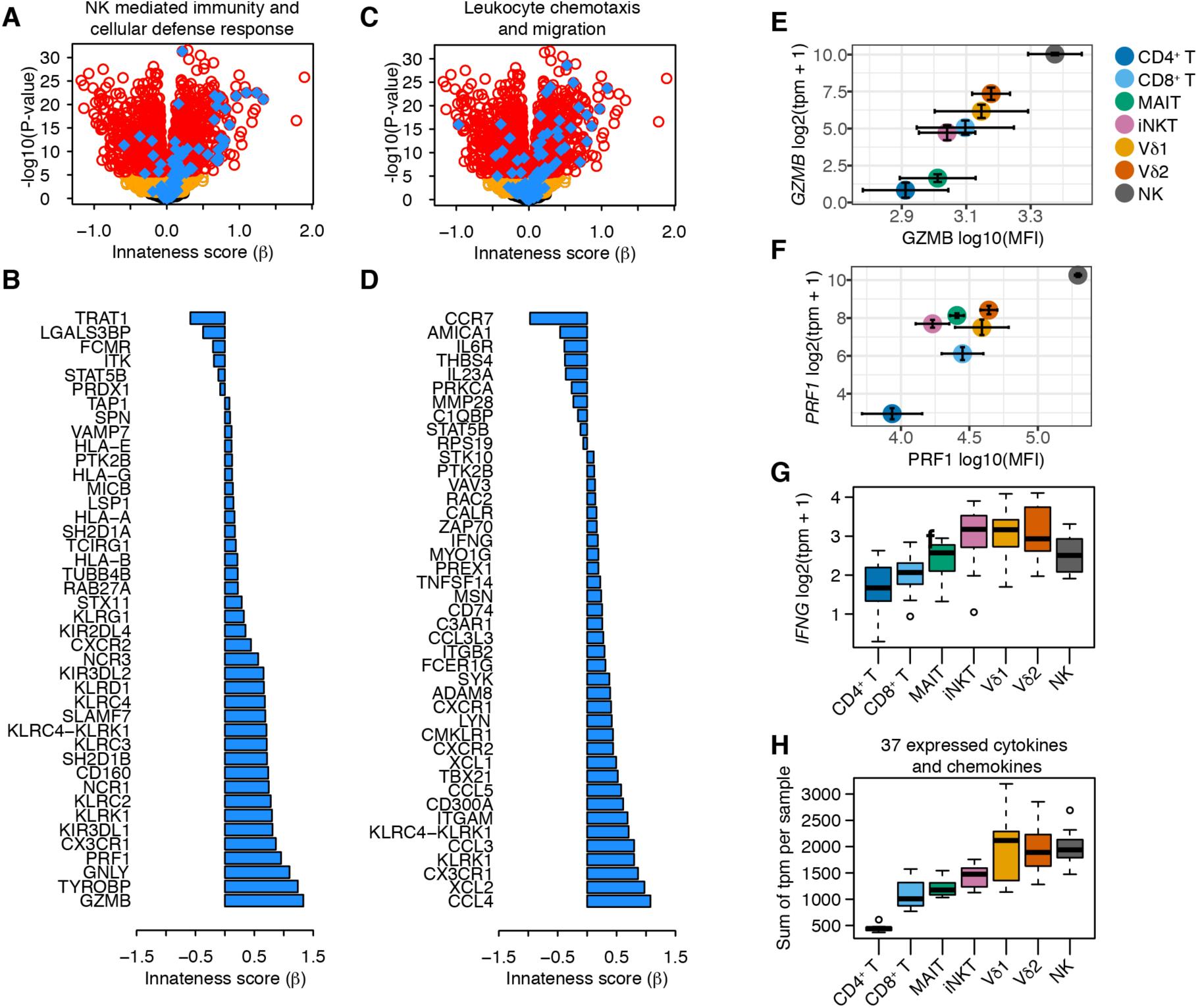
Genes and pathways associated with innateness. (A) Volcano plot showing associations with the innateness gradient. Yellow, genes with P < 0.05; red, genes with P < 2.5e-06 (Bonferroni threshold); blue, genes with GO terms involving NK mediated immunity and cellular defense response (GO:0006968, GO:0002228). (B) Innateness score (β) for individual genes in (A) with P < 2.5e-06. (C) As in (A) but with blue showing genes from GO terms involving leu ko cyte chemotaxis and migration (GO:2000501, GO:0035747, GO:1901623, GO:0030595, GO:2000401, GO:0097530, GO:0097529, GO:0048247, GO:0072676, GO:1990266). (D) Innateness score (p) for individual genes in (C) with P <25e-06. (E) GZMB and (F) PRF1 flow cytometric validation, showing protein levels (X-axis), and transcript levels with R NA-seq (Y-axis). (G) Distribution of IFNG transcript levels across all samples. (H) Sum of expression levels (tpm) for 37 cytokine and chemokine genes across all samples, per cell type. tpm, N=6; MFI, N=3. Error for tpm vs. MFI is s.e.m. Boxplots are described in methods.

#### Reactive oxygen species (ROS) metabolism

Metabolic pathways are well-known to vary among immune cell subsets and influence their functions (*27*). Among metabolic programs, the pentose phosphate pathway was nominally positively associated with innateness (P = 0.036, Fig. 4A). *G6PD*, the gene that codes for the rate-limiting enzyme in the pentose phosphate pathway, showed the strongest positive association with innateness in this pathway (*β* = 0.29, P = 3e-14, Fig. 4B,C). This enzyme produces NADPH which in turn can be used for glutathione biosynthesis, protecting against damage caused by ROS. Two critical enzymes for buffering the damaging effect of ROS, *GCLM* and *GCLC*, also nominally associated with innateness (P = 2e-04 and 1e-03, respectively, Fig. 4D,E). We quantified ROS by flow cytometry using CellROX green, and found that total cellular ROS levels were higher in adaptive T cells than in ITCs, suggesting that elevated *G6PD* might provide a baseline buffer counteracting ROS (Fig. 4F,G). Overall, these results suggest that ITCs are prepared to buffer ROS at baseline, a useful adaptation for effector cells expressing chemokine receptors such as CXCR1, CXCR2, and CCR5 (Fig. S7C) that direct them to the same sites of infection or inflammation as monocytes and neutrophils.

**Figure 4.**
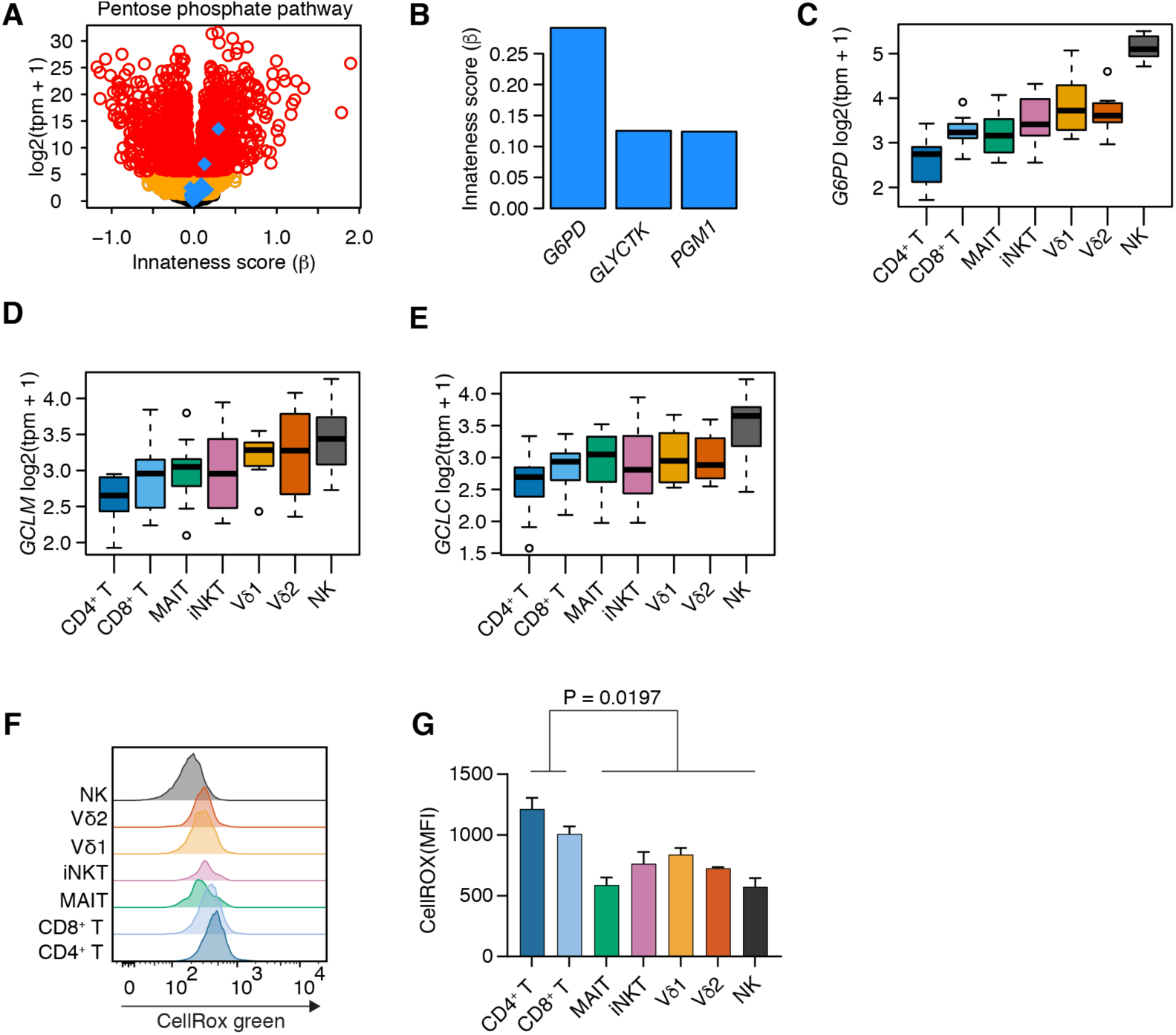
Elevated G6PD and lower ROS are associated with innateness. (A) Volcano plot showing associations with innateness gradient. Yellow, genes with P < 0.05; red, genes with P <2.5e-06 (Bonferroni threshold); blue, KEGG pentose phosphate pathway (hsa0003O). (B) Innate ness score (β) for genes in pentose phosphate pathway with P < 2.5e-06. Distribution by cell type for transcript levels of (C) G6PD, (D) GCLM, and (E) GCLC. (F) A representative plot and (G) N=3 with s.e.m., PBMCs labeled with CellRox green for one hour to quantify total cellular ROS, followed staining with markers to identify ITC populations. Boxplots are described in methods.

### Associations with adaptiveness: regulation of translational machinery

When we applied gene set enrichment to adaptiveness, “cytosolic ribosome” (GO:0022626) emerged as the most-associated term (P = 4.7e-28, Fig. 5A,B, Fig. S6A,B). This enrichment was not driven by a small percentage of genes very strongly overexpressed among ITCs (Fig. S6C). Translation initiation factors were also consistently associated with adaptiveness (Fig. 5C, Fig. S6D) suggesting that the translational machinery, and not just the ribosome complex, was associated with adaptiveness. *MYC*, which coordinately regulates ribosomal RNA genes (*28*), was the transcription factor with the highest fold change associated with adaptiveness (P = 3.8e-22, Fig. 5D). As an independent assessment of ribosome synthesis, we used quantitative polymerase chain reaction (qPCR) to assess expression of the earliest uncleaved ribosomal RNA (rRNA) precursor. The expression of precursor 47S rRNA associated with adaptiveness (Spearman rho = −0.57, P = 9e-05, Fig. 5E), suggesting that ITCs have a relative decrease in ribosome biogenesis.

**Figure 5.**
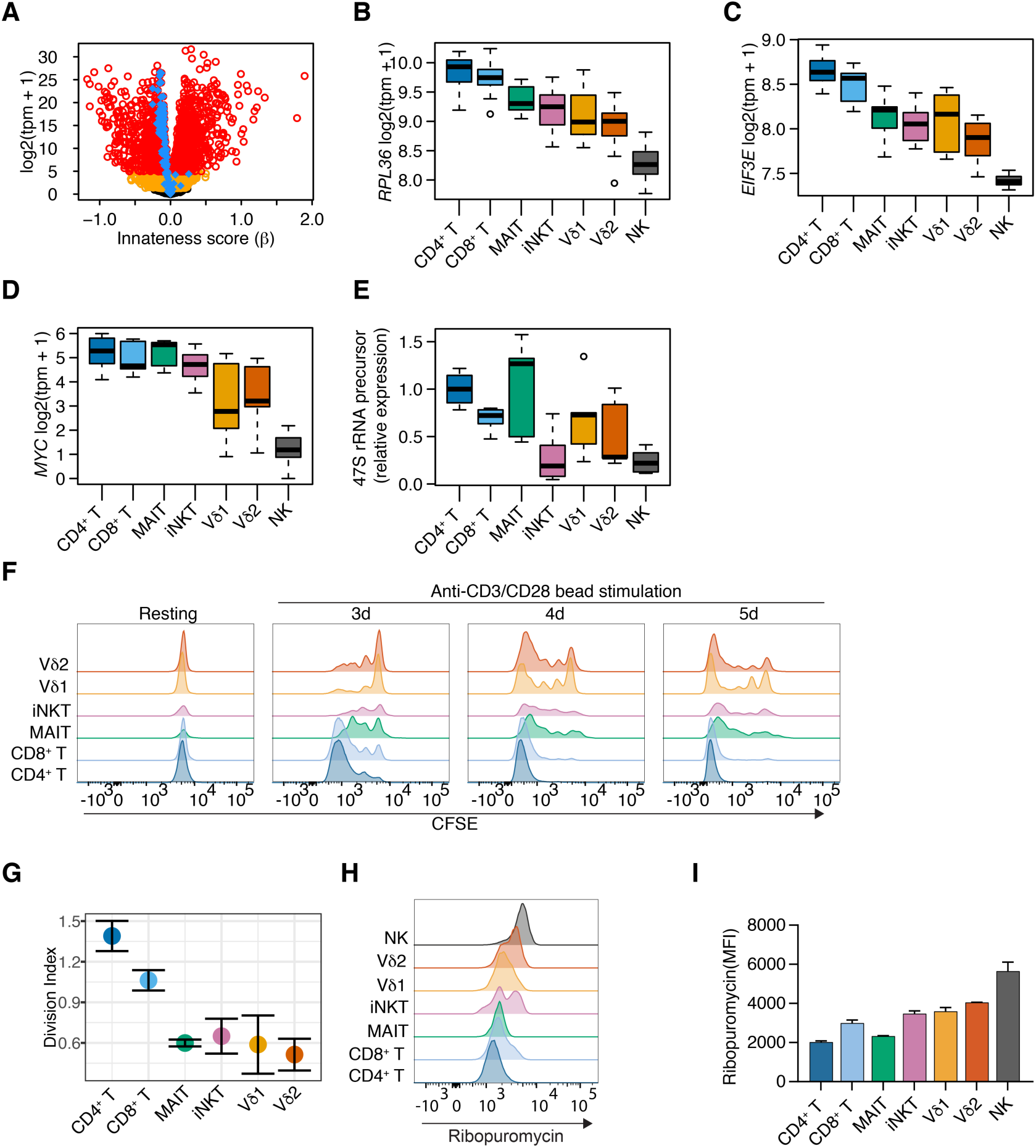
Reduced ribosome transcripts and proliferation in lTCs. (A) Volcano plot showing results of associations with the innateness gradient. Yellow, genes with P < 0.05; red, genes with P < 2.5e-06 (Bonferroni threshold); blue, GO term cytosolic ribosome (GO:0022626). Distribution by cell type for transcript levels of (B) ribosomal protein RPL36, (C) translation initiation factor EIF3E, and (D) MYC. (E) qPCR for 47S rRNA precursor from sorted cell populations (N =3). Expression shown is relative to HPRT, and normalized to CD4 T cells for comparison between samples. (F) PBMCs were labeled with CFSE then cultured with anti-CD3ICD28 coated beads before staining with markers to identify ITC populations, (G) division index shown at day 3 (N=3, s.e.m.). Ribopu romycylation in PBMCs to quantify total translational activity measured by flow cytometry, (H) representa tive plot and (I) N=3, s.e.m. Error for tpm vs. MFI is s.e.m. Boxplots are described in methods.

Since new ribosome production is necessary for proliferation, and *MYC* expression is generally associated with proliferative capacity, we hypothesized that proliferation potential might associate with adaptiveness. We found that proliferation in response to anti-CD3/CD28-coated beads, like *MYC* and ribosome biogenesis, associated with adaptiveness (Spearman rho = −0.73, P = 5.8e-04, Fig. 5F,G). We then assayed ribopuromycylation to quantify total active translation (*29*). Strikingly, we observed a positive association with innateness, with the innate cell types being engaged in more active translation than the adaptive T cells (Fig. 5H,I). This suggested that despite having lower expression of many major ribosomal genes, innate T cells have a higher number of ribosomes actively involved in translation. These results recall the well-described regulation of ribosomes in prokaryotes, where ribosome biogenesis is major energetic control point, is suppressed in conditions under which growth and division are deprioritized (*30*), and can be fine-tuned to ensure maximal occupancy of active ribosomes (*31*). Taken together, these results suggest that adaptive cells prioritize the production of factors required for cell growth and division, while innate cells may suppress ribosome biogenesis to prioritize the translation of other mRNAs, such as those encoding effector functions including the rapid production of cytokines (Fig. 1, Fig. 3G,H).

### Transcriptional regulation of innateness

We identified 142 transcription factors that varied significantly between cell types (F statistic, P < 5.8e-05, Bonferroni threshold). The expression of these transcription factors across cell types clustered into 4 major groups (Fig. 6A). Cluster 1 showed a gradual increase that closely matched the pattern of the innateness gradient. Cluster 2 showed a pattern opposite to that of cluster 1, with an increase in expression toward adaptive cellular populations. Cluster 3 showed high levels of expression in iNKT, MAIT, Vδ2, and NK cells, with relatively lower levels in adaptive T and Vδ1 T cells, and cluster 4 captured transcription factors with the opposite pattern to cluster 3 (Fig. 6A). In PCA of these transcription factors, the second principal component separated iNKT cells, MAIT, and Vδ2 T cells, from adaptive and Vδ1 T cells (Fig. S7A), similar to K-means clusters 3 and 4. These same cell groupings were also captured by PC2 generated using the overall most variable genes (Fig. 2A).

**Figure 6.**
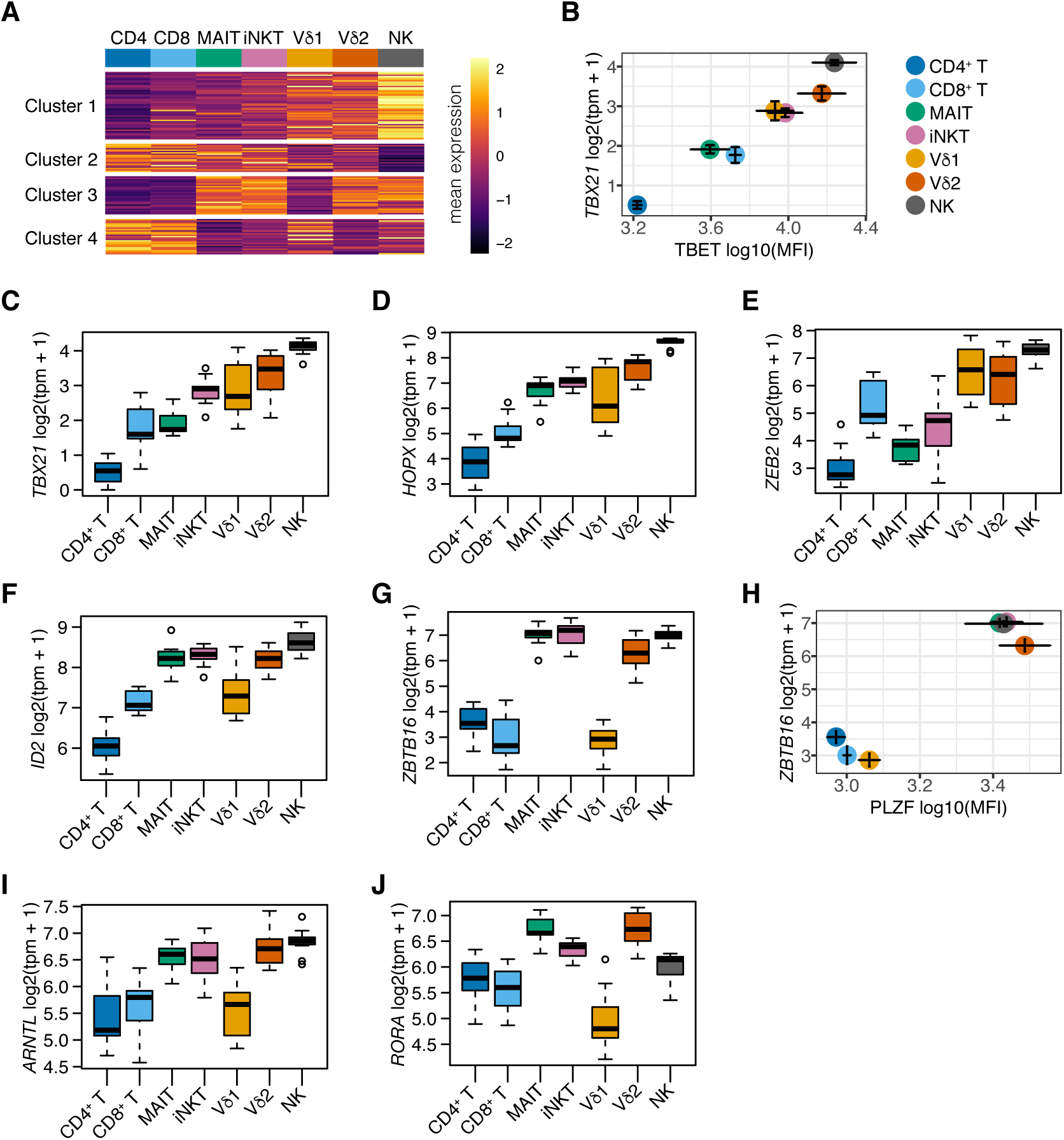
Transcription factors in ITCs. (A) Heatmap showing mean expression levels (scaled by row) for variable transcription factors among cell types, clustered into four groups. (B) Flow cytometric quantification of T-bet (N=3) compared to transcript levels with RNA-seq for its encoding gene TBX21 (N=6). Cluster 1, (C) TBX21, (D) HOPX, and (E) ZEB2; Cluster 3, (F ID2, (G) ZBTB16, (I) ARNTL, and (J) RORA. (H) Flow cytometric quantification of PLZF (N=3) compared to transcript levels for its encoding gene ZBTB16 (N=6). Error for tpm vs. MFI is s.e.m. Boxplots are described in methods.

Within cluster 1 of innateness-associated transcription factors, T-bet (*TBX21*, P = 2.4e-29), known for important roles in type 1 helper T cell (Th1), NK, and iNKT cell effector functions (*32-34*), followed the innateness gradient at both the transcript and protein levels (Fig. 6B,C). The next two innateness-associated transcription factors with the highest fold changes were *HOPX* and *ZEB2* (Fig. 6D,E). *HOPX* (P = 7.2e-25), reported to be induced by T-bet, has been shown to regulate persistence of effector memory Th1 cells, with upregulation in terminally differentiated cells (*35*). *ZEB2* (P = 1.8e-18) has been reported to cooperate with T-bet to induce terminal differentiation of cytotoxic T lymphocytes (*36, 37*). Two NFAT family proteins, *NFATC2* (P = 2e-16) and *NFAT5* (P = 1.1e-9), were associated with cluster 1 transcription factors. *IRF8* (P = 3.1e-14), *TFDP2* (P = 5.9e-13), *NFIL3* (P = 2.7e-13), *KLF10* (P = 2.1e-15), *RUNX3* (P = 5.6e-18), *LITAF* (P = 6.2e-24), *ZSCAN9* (P = 9.4e-17), and *ZNF600* (P = 4e-17) were also in this cluster and significantly associated with innateness. Within the adaptiveness-associated cluster 2, besides *MYC* (Fig. 5D), we found *TCF7*, involved in the maintenance of T cell identity (P = 2.4e-21) (*38*), *BACH2* (P = 4.3e-10), *NR3C2* (P = 1.8e-10), *POU6F1* (P = 1.9e-10), and *BCL11B* (P = 2e-17).

The third cluster of innateness-associated transcription factors, those enriched in iNKT cells, MAIT, Vδ2 T, and NK cells, included *ID2* (P = 8.8e-13, Fig 6F), *MYBL1* (P = 1.52e-10), *BHLHE40* (P = 8.1e-11), *FOSL2* (P = 8.8e-14), and *ZBTB16* (P = 1.1e-5, Fig 6G,H). Among these genes, *BHLHE40, FOSL2, ZBTB16* (encoding PLZF), and *ID2* have been reported to contribute to iNKT cell development and/or activation in mice (*39-43*). Id2 is also a major regulator of ILC development (*44*), and has been implicated Id2 in the regulation of mouse iNKT (*45*), ILC1 (*46*), and CD8^+^ T cell (*47*) effector functions in the periphery. Published transcriptional profiles of NK cells, ILC1, and influenza-specific Id2-deficient mouse CD8^+^ T cells showed a striking concordance of Id2-dependent expression with our innateness gradient genes, highlighted by *TBX21, ZEB2, IL18RAP, CCR7, TCF7*, cytotoxicity, and KLR genes (*46-48*). *TCF7*, consistently downregulated in ITCs, is negatively regulated by Id2, suggesting that in part, Id2 may drive the loss of adaptive T cell identity observed in ITCs. Taken together, these data suggest that Id2 may drive many features of innateness in human ITCs, and may be a major transcriptional node involved in maintaining their baseline innate state.

PLZF is a zinc finger transcription factor known to be important for the development and function of iNKT cells (*49, 50*), MAIT cells (*50*), and innate lymphoid cells (*51*). Mean PLZF protein expression by intranuclear staining confirmed our mRNA expression results (Fig. 6H). Human γδ T cells have previously been reported to express PLZF (*52*), but we did not detect elevated PLZF expression in Vδ1 cells (Fig. 6G,H). Differential expression analysis between PLZF^+^ ITCs and adaptive T cells revealed “cytokine receptor activity” as the most enriched term for upregulation in PLZF^+^ ITCs (P = 7.9e-05). PLZF expression in T cells was also associated with the aggregate expression of all cytokine and chemokine receptor activity genes (Fig. S7B), and we validated the expression of several of these receptors by flow cytometry (Fig. S7C). For genes differentially-expressed between PLZF^+^ ITCs and adaptive T cells, we found significant enrichment of PLZF target genes identified in mouse thymocytes with CHIP-seq (*53*) (P = 6.2e-07, Fig. S7D). In addition, PLZF^+^ ITCs upregulated genes that were associated with the term “circadian regulation of gene expression” (P = 4.2e-04), with major clock transcription factor genes like *ARNTL* (that codes for BMAL1), *RORA, PER1* and *CRY1* significantly upregulated in PLZF^+^ ITCs compared to adaptive T cells (P < 5e-08) (Fig. 6I,J, Fig. S7E). Both *BHLHE40* and *ID2* also have the capacity to regulate the circadian clock (*54-56*). Notably, although human NK cells express PLZF (mature mouse NK cells do not express PLZF), many genes upregulated in PLZF^+^ ITCs and identified as PLZF targets in mouse (*53*) showed low expression in human NK cells, including *CCR2, CCR7, CXCR6, RORC, CCR5, CCR6* and *LTK* (Fig. S7C,D). These results suggest that PLZF may regulate different sets of genes depending on the cell type, likely working as part of a larger gene network in determining ITC fate.

### Innateness in other populations of ITCs and adaptive T cells

We next investigated the innateness gradient in other candidate innate-like human T cell subsets. We chose two additional T cell populations for analysis, Vδ3-expressing γδ T cells and δ/αβ T cells, each of which can constitute up to 1% of human peripheral T cells (*57, 58*). We sorted Vδ3 T cells and δ*/αβ*T cells in duplicate from one individual and profiled their transcriptomes with ultra-low input RNA-seq. δ*/αβ* and Vδ3 clones have been identified that, like iNKT cells, recognize *α*-galactosylceramide presented by CD1d (*57, 58*), suggesting that these cells might potentially play a similar role in immunity to iNKT cells. However, principal component analysis revealed that δ*/αβ* T cells were closer to adaptive T cells, and closest to CD8^+^ T cells, rather than segregating with iNKT cells and other innate T cells (Fig. S8A,B). This suggests that δ*/αβ* T cells may have an adaptive-like phenotype. Vδ3 T cells, on the other hand, segregated closer to innate T cells by PCA, among the other γδ T cells (Fig. S8A,B). Neither δ*/αβ* T cells or Vδ3 T cells expressed PLZF.

Cytotoxicity genes and NK markers are expressed by a subset of adaptive T cells. We found that this class of genes was expressed by CD8^+^ T cells, and in some cases at higher levels than in ITCs. Interestingly, the development of innate-like Th1 effectors from adaptive cells has also recently been demonstrated in mice (*59*). To assess expression of innateness gradient genes in human adaptive effector T cells, we re-analyzed a human expression dataset generated using MHC class I tetramer-sorted, HCMV-specific CD8^+^ T cells (*60*) (polyclonal human CD8^+^ T cell datasets would likely be substantially ‘contaminated’ with ITCs). HCMV-specific effector memory CD8^+^ T cells expressed innateness gradient genes more highly than HCMV-specific memory CD8^+^ T cells (P = 1.4e-61, Wilcoxon paired test), which in turn had higher expression of these genes than naive CD8^+^ T cells (P = 7.9e-99, Fig. 7A). Conversely, genes associated with adaptiveness in our gradient were upregulated in naive CD8^+^ T cells compared to HCMV-specific memory CD8^+^ T cells (P = 9.2e-68), and also in memory CD8^+^ T cells compared to effector memory CD8^+^ T cells (P = 2.7e-11, Fig. 7A). We also re-analyzed published RNA-seq data for CD4^+^ T cell subsets (*61*). CD4^+^ effector memory T cells had higher expression of innateness-associated genes than CD4^+^ naive and CD4^+^ central memory T cells (P < 6.4e-119, Fig. 7B), whereas naive CD4^+^ T cells had higher expression of adaptiveness-associated genes than CD4^+^ central memory and effector memory T cells (P < 1.4e-54, Fig. 7B). Overall, these results suggest that the innateness gradient can stratify adaptive effector CD8^+^ and CD4^+^ T cell populations following infection, and is thus not limited to ITCs.

**Figure 7.**
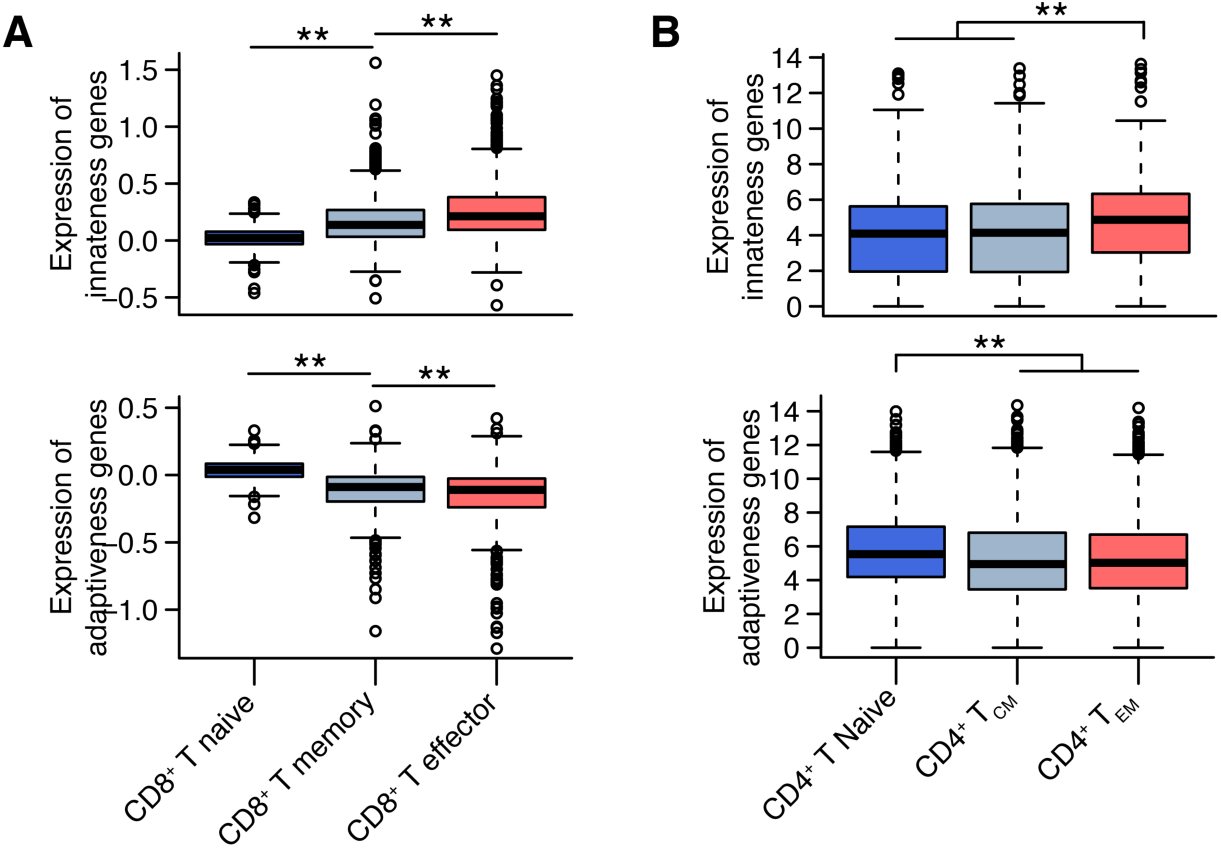
Innateness gradient genes in other T cell populations. Re-analysis of innateness gradient genes in (A) HCMV-specific CD8 T cell populations (naive =CD45RA^+^CD27bright; memory CD45RA^−^CD27^+^; effector CD45RA^+^CD27), and (B) CD4^+^ T cell popula tions (naive = CCR7^+^CD45RA^+^CD45RO^−^; T central memory (T_CM_) = CCR7^+^CD45RA^−^CD45RO^+^; T effector memory (T_EM_) = CCR7^−^CD45RA^−^CD45RO^+^). Depicted are distributions of mean expression levels for innateness genes (upper) and adaptiveness genes (lower), averaged across 4 replicates for (A) and 5 replicates for (B). ^**^ P < 3e-11. Boxplots are described in methods.

## Discussion

MAIT, iNKT, γδ, and other innate-like T cells do not fit neatly into traditional paradigms of adaptive or innate immunity. Their nature has been an interesting puzzle for more than 30 years. Each population has been studied in depth individually, but rarely have they been considered in aggregate. Here, we set out to study human ITCs as a group, addressing two important questions, 1) is there a shared transcriptional basis for their functions in immunity, and 2) how do ITCs maintain their baseline effector state? In quantitative, unbiased analyses, we discovered that ITCs segregate along an innateness gradient between prototypical adaptive and innate populations. We propose that the large transcriptional programs positively-and negatively-associated with this gradient represent the transcriptional basis of lymphocyte innateness. Our data support that ITCs are indeed a ‘family’ with a common transcriptional basis for their similar functions in immunity, including rapid cytokine and chemokine production, chemotaxis to areas of inflammation, cytotoxicity, and TCR-independent responses. The functional and transcriptional conservation of innate-like functions in ITCs suggests that they enhance evolutionary fitness. That humans dedicate such a large part of their T cell repertoire to the generation of innate-like receptors is a testament to the teleological importance of innate immune surveillance even after the evolution of adaptive immunity.

Strikingly, we observed that this innateness program can not only classify ITCs according to their innateness, but can also differentiate adaptive effector populations. For example, naive, memory, and effector adaptive populations can be separated by their innateness. Interestingly, both Th1 and Th2 adaptive T cells have been demonstrated to acquire innate-like characteristics in some settings (*62-65*). Thus, the study of ITCs highlights important pathways used across innate and adaptive lymphocyte populations.

The shared gene programs associated with innateness included cytokine/chemokine production, cytotoxicity, and cytokine/chemokine receptor expression. For the genes positively associated with the innateness gradient, this is essentially an ‘effector gradient,’ which strongly supports a role for ITCs in host defense. We found that human ITCs rapidly produced IFN-*γ* after activation through their TCRs (Fig. S2), as do a smaller fraction of adaptive T cells. However, IFN-*γ* production in response to IL-12, IL-18, and IFN-*α*, cytokines generated by myeloid or stromal cells in response to danger signals, were almost exclusively limited to ITCs (Fig. 1B). This is consistent with the role of ITCs as innate responders where prior pathogen experience is not required. Thus, T cell innateness can regulate the response to pathogen-associated molecular patterns. Of note, human Vδ1 cells have been demonstrated to be variable in both TCR repertoire and numbers, and likely respond to specific infections (*66*). Although they express much of the innateness program, Vδ1 cells may not fit the ITC paradigm as neatly as the more-conserved MAIT, iNKT, and Vδ2 populations. Indeed, Vδ1 cells have greater TCR diversity, exhibit less ‘cytokine-only’ activation, and do not express PLZF.

Our identification of ribosome subunits and other factors involved in translational activity associated with adaptiveness (Fig. 5) also sheds light on the biology of ITCs. For an adaptive T cell, population expansion is of central importance in both primary and recall immune responses. ITCs, on the other hand, are likely to function as sentinels early during infection, acting as ‘cellular adjuvants’ to enhance the larger immune response in response to microbial molecules. For such a role, rapid effector responses are key, and proliferation may serve only to replenish numbers at a later stage. Taken together, the effector-focused transcriptional programs of ITCs and proliferation-focused programs of adaptive cells are ideally suited to support their respective roles in immunity.

Finally, the innateness gradient reported here could be applied in different scenarios in order to better understand human immunology. A transcriptomic innateness score could be employed as a unified T cell metric to classify individual single cells assayed with single-cell RNA-seq, and could provide a better understanding of patient heterogeneity. We can use our immunoprofiling data and create an ‘innateness metric’ for each individual based on the abundance of each T cell type weighted by the innateness level of that cell type. This score is remarkably variable between individuals, even after correcting for age (Fig. S9). This single innateness metric in an individual might be associated with genetic differences, human diseases including cancer, infection, and allergy, or therapeutic responses to immunomodulating medications.

## Materials and Methods

### Study design

To study the transcriptome of innate T cell populations (MAIT, iNKT, Vδ1, Vδ2), we compared them with adaptive cells (CD4^+^ T, CD8^+^ T) as well as NK cells as prototypical innate lymphocytes. Samples used for immunophenotyping and RNA-seq analyses were from healthy individuals. All human sample use was approved by the Brigham and Women’s Hospital Institutional Review Board, including direct consent for public deposition of RNA sequencing. A matched set of populations were sorted from each individual to avoid batch effects. All blood draws were performed in the morning, and cells were immediately stained and double-sorted directly into lysis buffer. Based on previous RNA-seq analyses on number of replicates and read depth for optimal differential expression analysis (*67*), we decided to sort cells from 6 individuals in duplicate (total of 12 samples per cell-type) at a read depth of 4-12 million read pairs (8-24 million reads). The goal of this study was to define the shared transcriptional programs between cell populations rather than variability between individuals. To avoid systematic technical error or batch effects, samples were randomized within the plate for library preparation, and all samples sequenced together. Five samples were removed for low read depth (described below).

### RNA library preparation and sequencing

Smart-seq2 libraries (*68*) (poly-A selected) were prepared for the 90 flow-sorted samples (each 1,000 cells). These samples were composed of 7 main cell types (CD4^+^ T, CD8^+^ T, MAIT, iNKT, Vδ1, Vδ2 and NK cells) from 6 healthy donors, and 3 additional cell types (δ*/αβ*,Vδ3 and B NK cells) from one healthy donor. Each sample had 2 duplicates. Samples were randomized within plate. 25 base paired-end sequencing was performed yielding 4-12M read pairs (8-24e6 reads, Fig. S4)

### Gene expression quantification

We used Kallisto version 0.43.1 (*69*) to quantify gene expression using the Ensembl 83 annotation. We included protein-coding genes, pseudogenes, and lncRNA genes. As expected, protein coding genes were the most highly expressed, followed by lncRNAs and then pseudogenes (Fig. S4C). We removed 5 outlier samples that had low proportion of common genes detected (1 MAIT, 1 CD8^+^ T, 1 NK, and two Vδ1 samples; Fig. S4D). We used log-transformed tpm (transcripts per million) as our main expression measure, which accounts for library size and gene size (specifically log2(tpm+1)). We considered as expressed genes those with a log2(tpm+1) > 2 in at least 10 samples. We further performed quantile normalization on the log2(tpm+1) values for our differential expression analyses. Boxplots were created in R. Boxes show the 1^st^ to 3^rd^ quartile with median, whiskers encompass 1.5X the interquartile range, and data beyond that threshold indicated as outliers.

### Differential expression analyses

We used linear mixed models for our differential expression and expression association analyses. The dependent variable was quantile normalized log2(tpm+1) expression values. Within our predictor variables, we used in all cases donor ID as a random effect. For associations with the innateness gradient, we used one fixed effect composed of integers from 1-7 (for CD4^+^ T, CD8^+^ T, MAIT, NKT, Vδ1, Vδ3 and NK, respectively). In the differential expression between adaptive cells and PLZF^+^ ITCs we used one fixed effect taking values of 0 or 1, respectively.

### Gene ontology term enrichment analyses

We downloaded Ensembl gene IDs linked to Gene Ontology (GO) terms on April 2016 (*70, 71*). This included 9,797 GO terms and 15,693 genes. We tested for GO enrichment sorting genes by the *β* (effect size) of our differential expression analysis. We used the minimal hypergeometric test (*72*) to test for significance. We confirmed significance of enrichment for the top GO terms using an alternative method: the function gsea of the liger package (https://github.com/JEFworks/liger).

### Pathway enrichment analysis

We downloaded genes pertaining to 12 KEGG pathways (*73*) from the Consensus Pathway Database-human http://cpdb.molgen.mpg.de/ (*74*) in March 2017. First, we calculated the F statistic per expressed gene in our dataset as a metric of variability between cell types. Then we tested whether the F statistics in genes of a certain pathway were higher than the other expressed genes using a Wilcoxon test. Three pathways had a P-value < 0.05. Since higher expressed genes tend to have higher F statistics, we further tested whether these 3 pathways had significantly higher F statistics than expected by controlling for gene expression. Specifically, we chose a null set of genes with similar expression levels by taking for each gene in a pathway, 30 random genes with mean level of expression (across all cell types) within 10% of the standard deviation. After this, only the pentose phosphate pathway had genes with F statistics higher than expected (P = 0.018). We further tested enrichment of this pathway in genes associated with innateness gradient using the gsea function of the liger package (Fig. 4A).

### Immunophenotyping associations

Associations among cell types and clinical traits, when accounting for different covariates, were tested with linear regression using cell type percentages in log scale. For iNKT cell abundance, there were 2 individuals with zero values, and these were converted to the next minimal value of 0.01 before log transformation.

### PLZF target analysis

We downloaded PLZF ChIP-seq peaks from the Gene Expression Omnibus (GEO) database from Mao et al (*53*) (accession number GSE81772). We used genes from the mouse Gencode vM14 annotation. We defined gene targets as mouse genes with a PLZF peak in the gene body or within 2kb from the transcription start site (TSS). We downloaded mouse-human gene homologues from BioMart (*75*). We selected only genes with 1 to 1 orthologues. We then checked from the mouse PLZF gene targets to which human orthologue they correspond. Finally, we performed logistic regression to determine whether gene targets are enriched in differentially expressed genes between PLZF^+^ ITCs and adaptive T cells. Specifically, the response variable is 0 or 1 for non-target or target gene, respectively. The predictor variable was the *β* of the differential expression analysis of PLZF^+^ ITCs versus adaptive T cells. We also tested enrichment defining gene targets if a peak was found only at the promoter region of a gene (−2kb to +1kb from TSS), and found similar results.

### Data accessibility

Our RNA-seq data is available at GEO with accession number TBD. Processed expression data and innateness gradient associations can be viewed using an interactive browser at https://immunogenomics.io/itc.

### Flow cytometry and cell sorting

For immunophenotyping, Ficoll-isolated (GE Healthcare) PBMCs were prepared within 2 hours of overnight fasting with blood draw between 8 and 10 AM, stained, and data was acquired the same day. For sorting, freshly-isolated PBMCs from donors that had at least 0.1% for each cell type were processed in accordance with the ImmGen standard operating procedure (*76, 77*). Briefly, after Fc receptor binding inhibitor (eBioscience), cells were stained with surface antibodies and dead cells identified with 7-AAD (Biolegend). Using a FACSAria Fusion sorter fitted with a 100 µM nozzle, 1,000 cells double-sorted in duplicate directly V-bottom plates with TCL lysis buffer (Qiagen) and stored frozen until processing.

For validation studies, cryopreserved PBMCs were used from a total of 15 donors. The antibodies used for flow-cytometric validation are listed separately. Data was acquired with a 5-laser LSR Fortessa or 3-laser FACSCanto II (BD Biosciences) and analyzed with FlowJo (Treestar). A live-dead dye was used for all staining, either 455UV (eBioscience) or ZombieAqua (Biolegend). The gating strategy for these studies is shown in Fig. S3. For intracellular cytokine production studies, cells were fixed with 4% paraformaldehyde, then permeabilized with BD Perm/Wash (BD Biosciences) and stained with intracellular antibodies. For intranuclear staining to assess expression of transcription factors, cells were fixed and permeabilized using the FoxP3 buffer set (eBioscience). For validation studies, MAIT cells were identified as V*α*7.2^+^CD161^+^ T cells.

### qPCR analysis

For 47S rRNA quantification, cells were sorted directly into RLT buffer (Qiagen) before RNA extraction (Qiagen, RNeasy). Primers were designed to span the first rRNA processing site using the following sequences: forward: GTCAGGCGTTCTCGTCTC, reverse: GCACGACGTCACCACAT. HPRT was used as a housekeeping control (forward: CGAGATGTGATGAAGGAGATGG, reverse: TTGATGTAATCCAGCAGGTCAG). qPCR was performed using the Brilliant III Ultra-Fast SYBR QPCR Master Mix (Agilent), read on a Stratagene MX3000P system.

### Cell culture, activation, and proliferation studies

For cellular activation studies, PBMCs were cultured in RPMI 1640 supplemented with 10% FBS (Gemini), HEPES, penicillin/streptomycin, L-glutamine, and 2-mercaptoethanol. Cytokines were from Peprotech except for IFN-*α* (R&D Systems). For assessment of cytokine production, PMA (200 ng/ml, Sigma) and Ionomycin (500 ng/ml, Sigma) were added along with Protein Transport Inhibitor Cocktail (eBioscience) containing brefeldin and monensin for 4 hrs. Cytokine production in response to IL-12 (20 ng/ml), IL-18 (50 ng/ml), and IFN-*α* (50 ng/ml), PBMCs were cultured for 16hrs with these cytokines, with eBioscience Protein Transport Inhibitor Cocktail added for the last 4 hrs of culture. For measurement of cellular ROS, PBMCs were thawed, rested overnight in complete media without added cytokines, followed by the addition of CellRox Green (Thermo Fisher) for 1 hr. For proliferation, cells were labeled with CFSE (5 *μ*M for 5 min in PBS), then cultured at a 2:1 ratio with anti-CD3/CD28-coated beads (Dynabeads, Thermo Fisher). Division index was calculated as (cells divided once + (cells divided twice/2) + (cells divided ≥ 3 times / 2.67)) / (undivided cells + (cells divided once/2) + (cells divided twice/4) + (cells divided ≥ 3 times / 8)) (FlowJo, TreeStar).

### Ribopuromycylation studies

To assess ribosomal activity, we adapted a microscopic technique, ribopuromycylation (*29*) for use by flow cytometry. Puromycin was added for 5 min in the presence of emetine (100 *μ*g/ml), followed by fixation with 4% paraformaldehyde, permeabilization with BD Perm/Wash, and staining with an antibody that recognizes puromycin (EMD Millipore).

### List of antibodies

| **Name** | **Clone** | **Company** |
| --- | --- | --- |
| CCR1 | 5F10B29 | Biolegend |
| CCR2 | K036C2 | BioLegend |
| CCR5 | HEK/1/85a | BioLegend |
| CCR6 | G034E3 | BioLegend |
| CCR7 | G043H7 | BioLegend |
| CD117 | 104D2 | eBioscience |
| CD161 | HP-3G10 | BioLegend |
| CD1d-PBS57 tetramer | NA | NIH tetramer facility |
| CD3 | UCHT1 | eBioscience |
| CD4 | RPA-T4 | BioLegend |
| CD56 | 5.1H11 | BioLegend |
| CD8a | RPA-T8 | BioLegend |
| CellRox Green | NA | Thermo Fisher |
| CXCR6 | K041E5 | BioLegend |
| DAP12 | 406288 | R&D Systems |
| EOMES | WD1928 | eBioscience |
| GATA3 | 16E10A23 | BioLegend |
| Granulysin | DH2 | BioLegend |
| Granzyme A | CB9 | eBioscience |
| Granzyme B | GB11 | eBioscience |
| Granzyme K | G3H69 | eBioscience |
| IFNγ | B27 | BioLegend |
| IL18Ra | H44 | eBioscience |
| KIR2DL1 | HP-MA4 | BioLegend |
| KIR2DL3 | DX29 | BioLegend |
| LTK | AF4664 (sheep polyclonal) | R&D Systems |
| MR1-5-OP-RU tetramer | NA | gift from J. McCluskey |
| NKG2D | 1D11 | eBioscience |
| NKp46 | 9E2 | Biolegend |
| NKp80 | 5F12 | BioLegend |
| Pan $\gamma\delta$ TCR | B1 | BioLegend |
| Perforin | B-D48 | Biolegend |
| PLZF | R17-809 | BD Pharmingen |
| Puromycin | 12D10 | EMD Millipore Corp |
| RORC | AFKJS-9 | eBioscience |
| T-bet | eBio4B10 | eBioscience |
| TCR Vα7.2 | 3C10 | BioLegend |
| TCR $\alpha\beta$ | T10B9.1A-31 | BD Pharmingen |
| TCRδ (to identify $\delta/\alpha\beta$) | A13 | gift from L. Moretta |
| TIGIT | A1513G | BioLegend |
| Vδ1 TCR | REA173 | Miltenyi Biotec |
| Vδ2 TCR | B6 | BioLegend |
| Vδ3 TCR | P11.5B | gift from L. Lynch |

## Supplementary Materials

Fig. S1. Innate T cell population frequency associations with age and covariance.

Fig. S2. Pharmacologic and cytokine-only activation of lymphocyte populations.

Fig. S3. Gating strategy used for sorting for RNA-seq and validation experiments.

Fig. S4. RNA-seq summary and quality check.

Fig. S5. Innateness-associated genes and pathways.

Fig. S6. Ribosome associations with adaptiveness.

Fig. S7. Transcription factors in ITCs.

Fig. S8. Innateness in candidate ITCs and adaptive T cells.

Fig. S9. Individual innateness metric.

Table S1. T cell subset abundances and clinical information of 101 individuals.

Table S2. T cell subset samples isolated and RNA sequenced.

Table S3. Results of gene expression associations with innateness gradient.

## Acknowledgements

We thank J. McCluskey, L. Morretta, L. Lynch, the National Institutes of Health Tetramer Facility, and the Kraft Family Blood Donor Center for providing critical materials. We thank members of the Brennan, Raychaudhuri, and Brenner laboratories, as well as S. Suliman, P. Ivanov, S. Lyons, N. Kedersha, and M. Fay for thoughtful discussions and/or critical reading of the manuscript. We thank S. Pathak for facilitating pilot data. Supported by the National Institutes of Health (AI102945 to P.J.B, AI113046 and AI063428 to M.B.B, U19AI111224, U01GM092691, U01HG009379 and R01AR063759 to S.R.), the Doris Duke Charitable Foundation (2013097 to S.R.), the Violin and Karol families (to P.J.B), the Swiss National Science Foundation (Early Postdoc Mobility Fellowship to M.G.-A.).

## Supplementary Materials

**Supplementary Figure 1.**
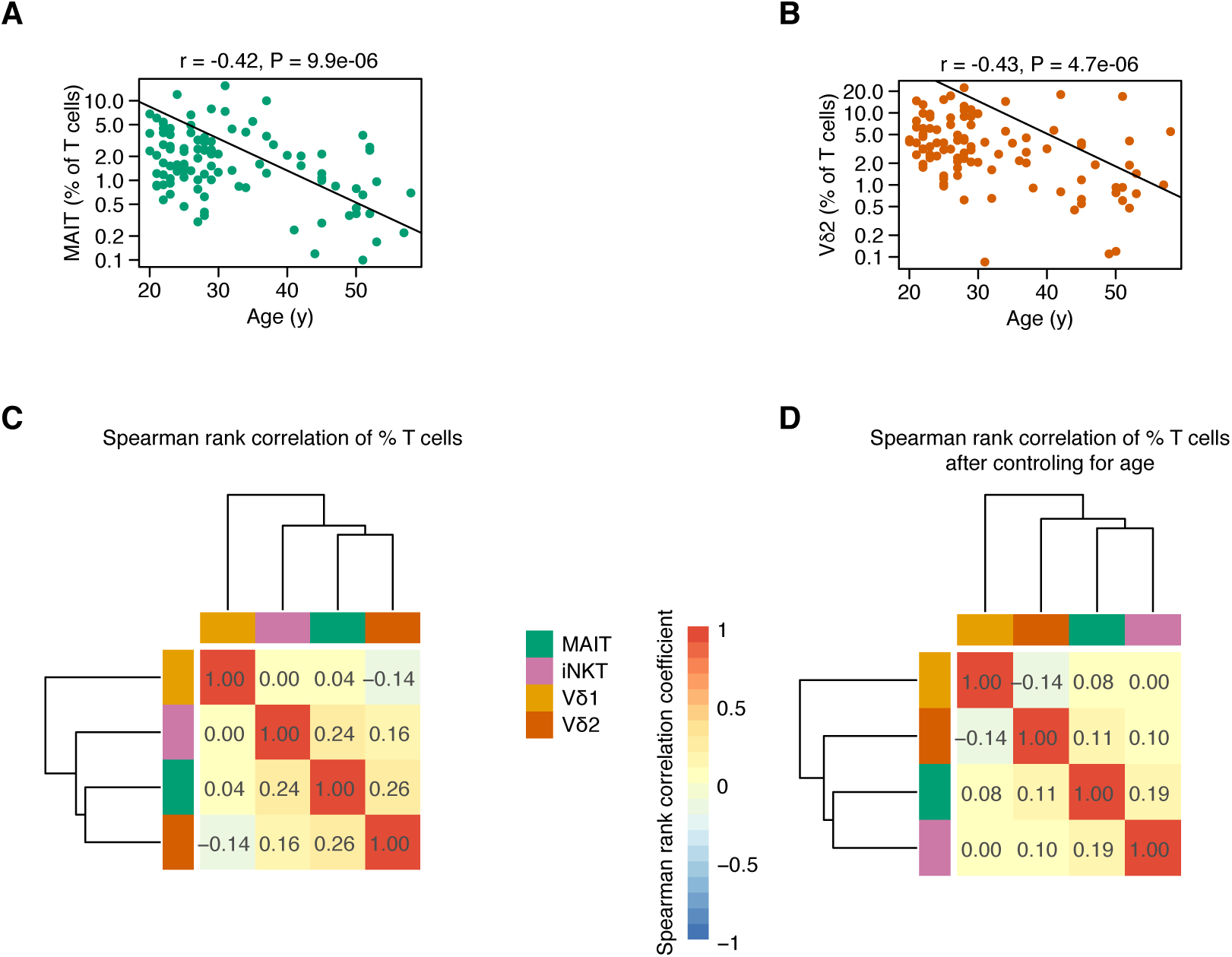
Innate T cell population frequency associations with age and covariance. Associations between donor age and (A) MAIT or (B) Vδ2 T cells, P-value and r from Pearson correla-tions. Heatmaps depict pairwise Spearman correlation coefficients between T cell count percentages in 101 healthy individuals, (C) before and (D) after regressing out age effects.

**Supplementary Figure 2.**
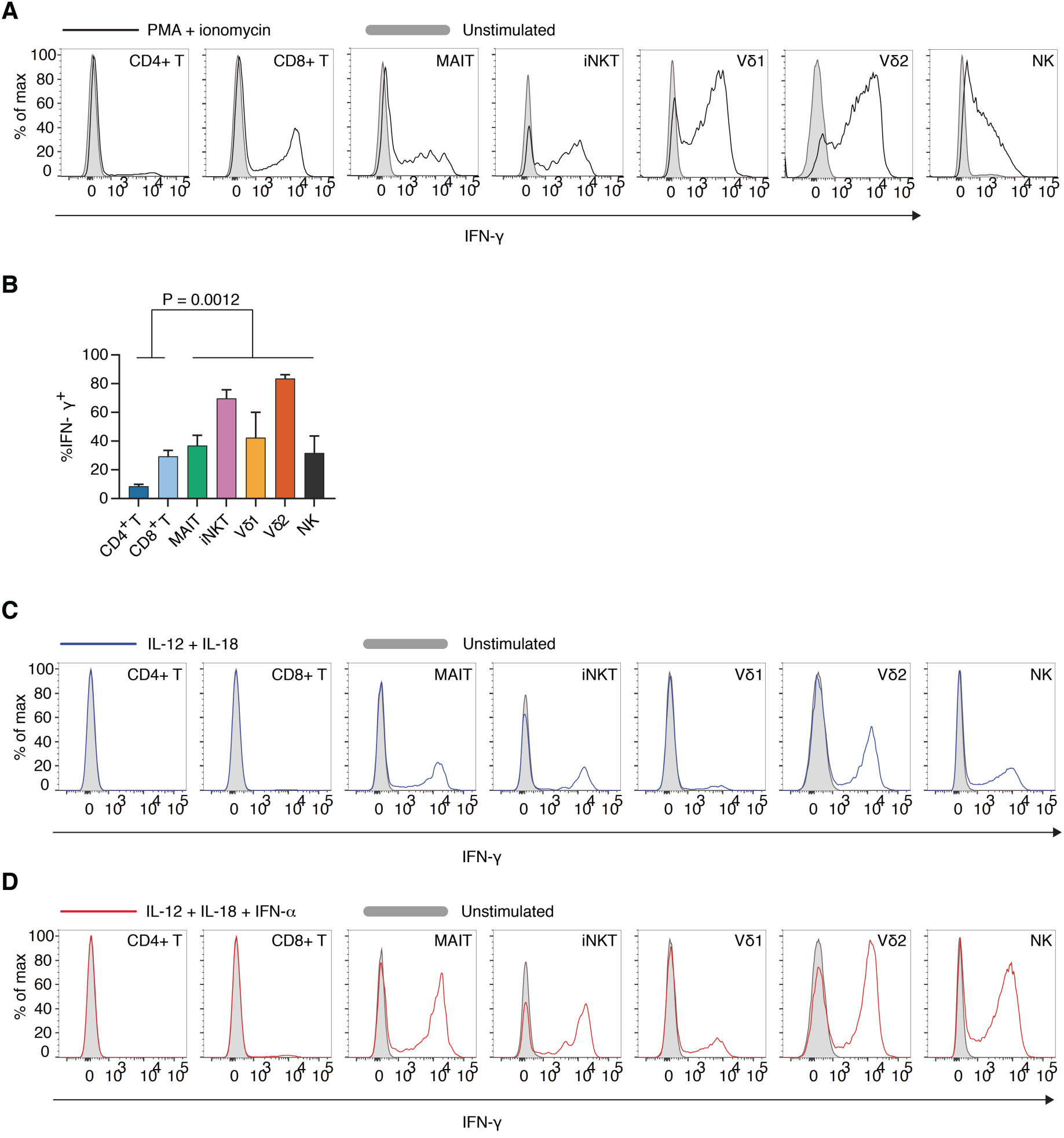
Pharmacologic and cytokine-only activation of lymphocyte populations. PBMC were activated with PMA and ionomycin for 5 hrs and IFN-y production was quantified by intra cellular cytokine staining, (A) a representative plot and (B) N=3, SEM. Representative flow plots for PBMC activated for 16 hrs with (C) IL-12 and IL-18 or (D) IL-12 + IL-18 + IFN-α, with lEN-γ production quantified during the last 4 hrs by intracellular cytokine staining.

**Supplementary Figure 3.**
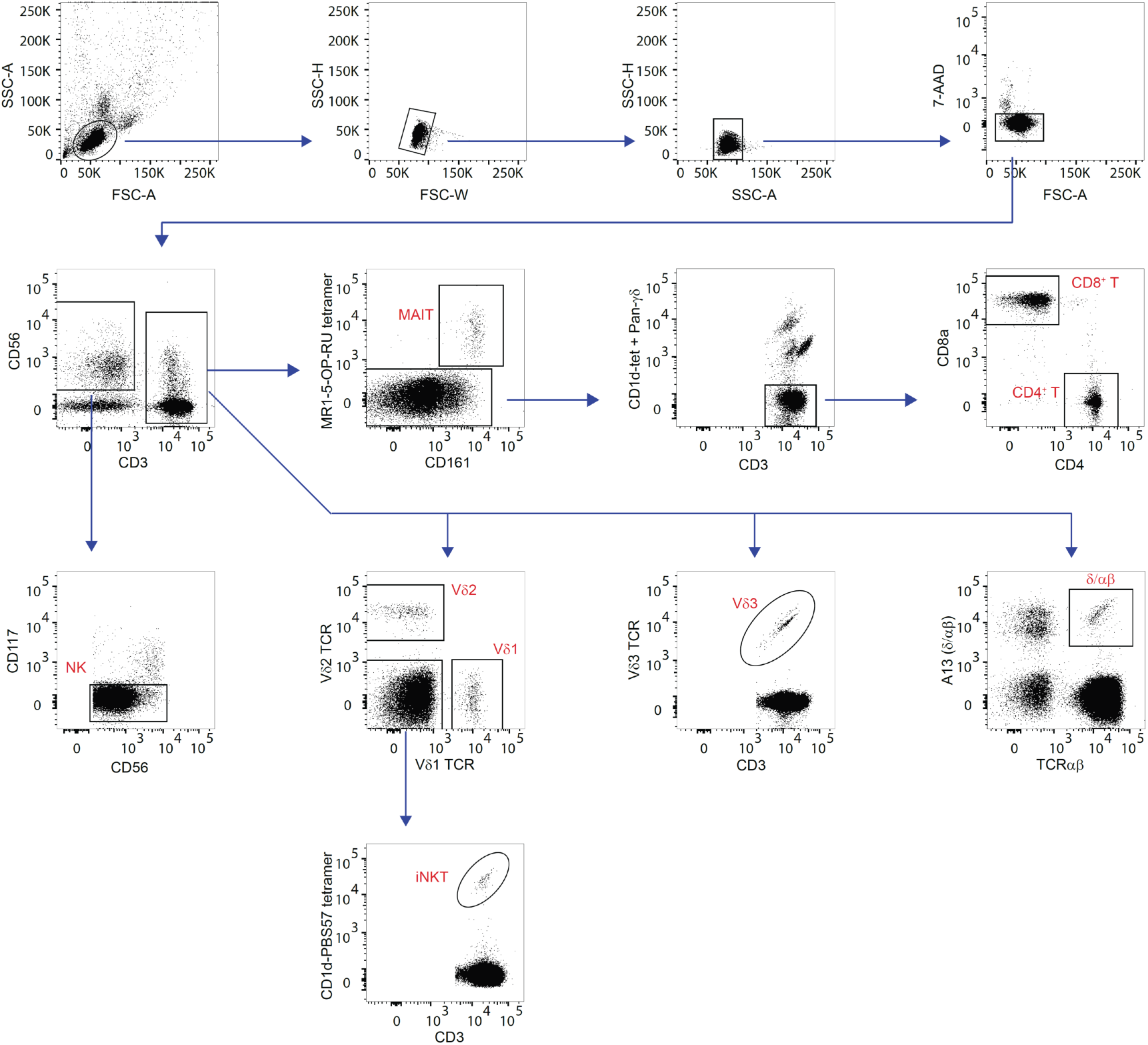
Gating strategy used for sorting for RNA-seq and validation experiments. For validation studies, MR1-5-OP-RU tetramer was replaced by anti-Vα7.2 TCR.

**Supplementary Figure 4.**
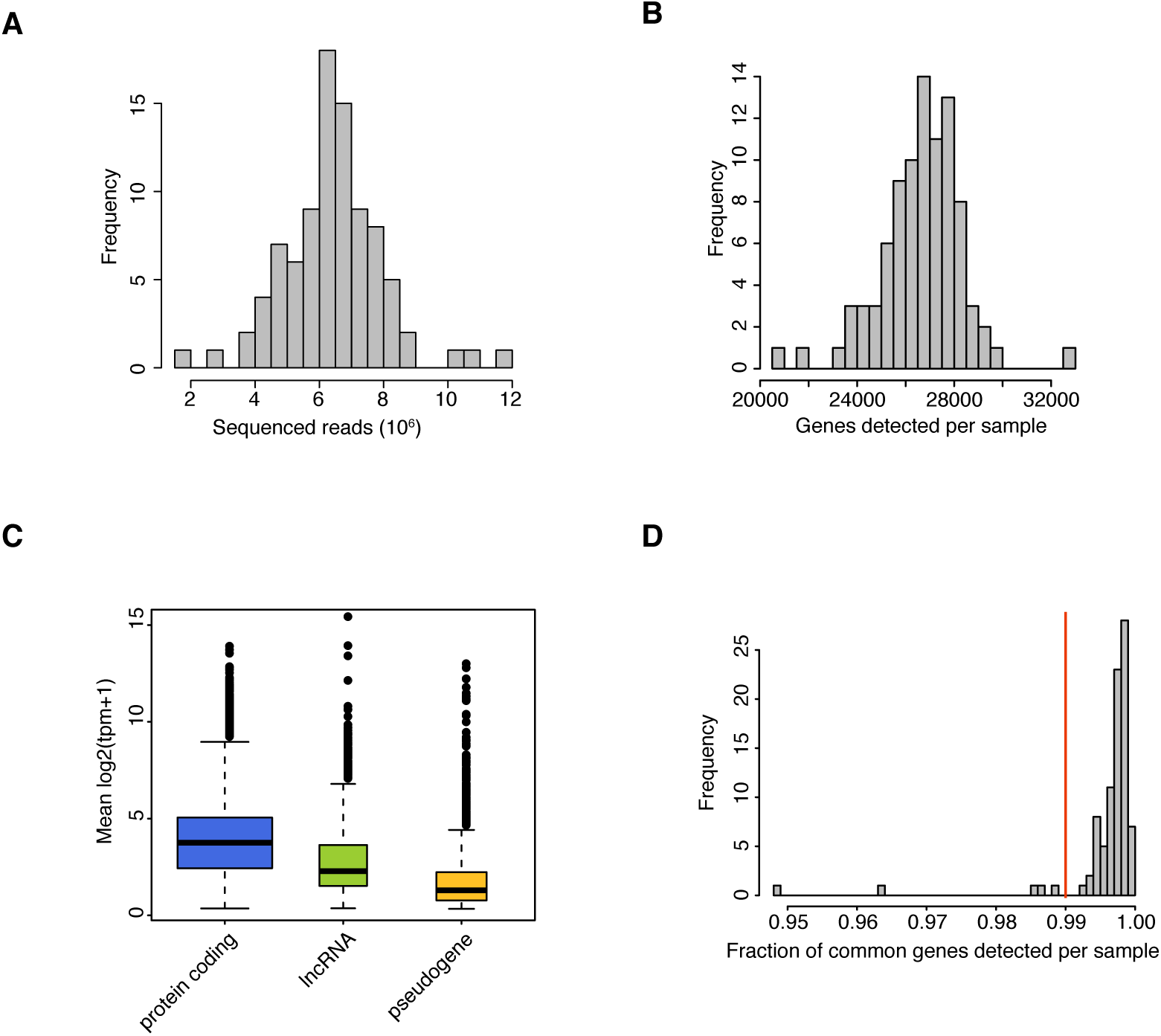
RNA-seq summary and quality check. (A) Number of sequenced reads, and (B) number of genes detected per sample. (C) Distributions of mean expression levels for protein-coding genes, lncRNA genes, and pseudogenes. (D) Fraction of common genes detected per sample. Samples to the left of the vertical red line were considered low quality and were discarded from further analyses.

**Supplementary Figure 5.**
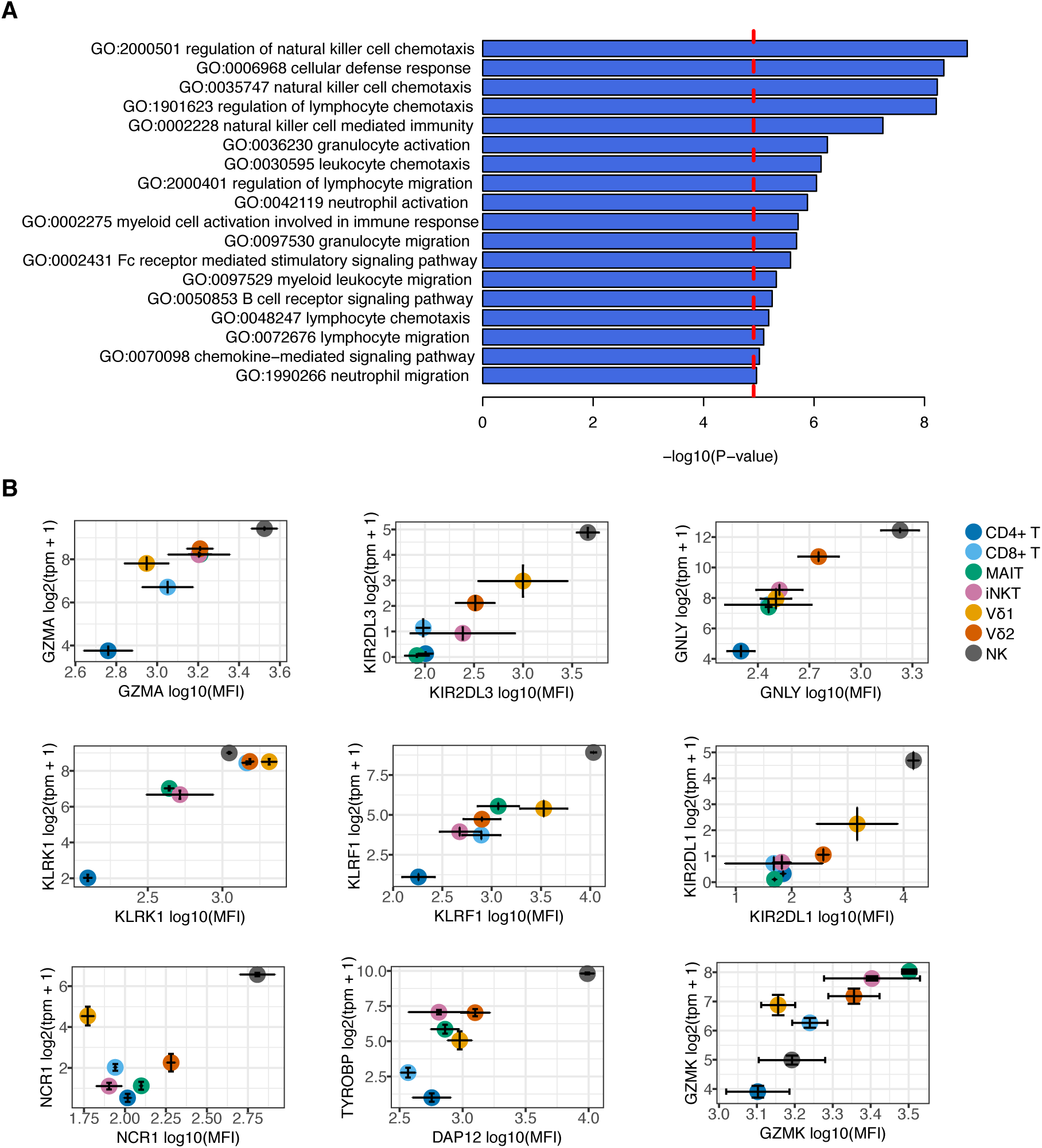
Innateness-associated genes and pathways. (A) GO terms significantly associated within innateness genes. (B) Flow cytometric validation for innateness associated genes, showing protein levels by flow cytometry (X-axis), and transcript levels with RNA-seq (Y-axis). tpm, N=6; MFI, N=3.

**Supplementary Figure 6.**
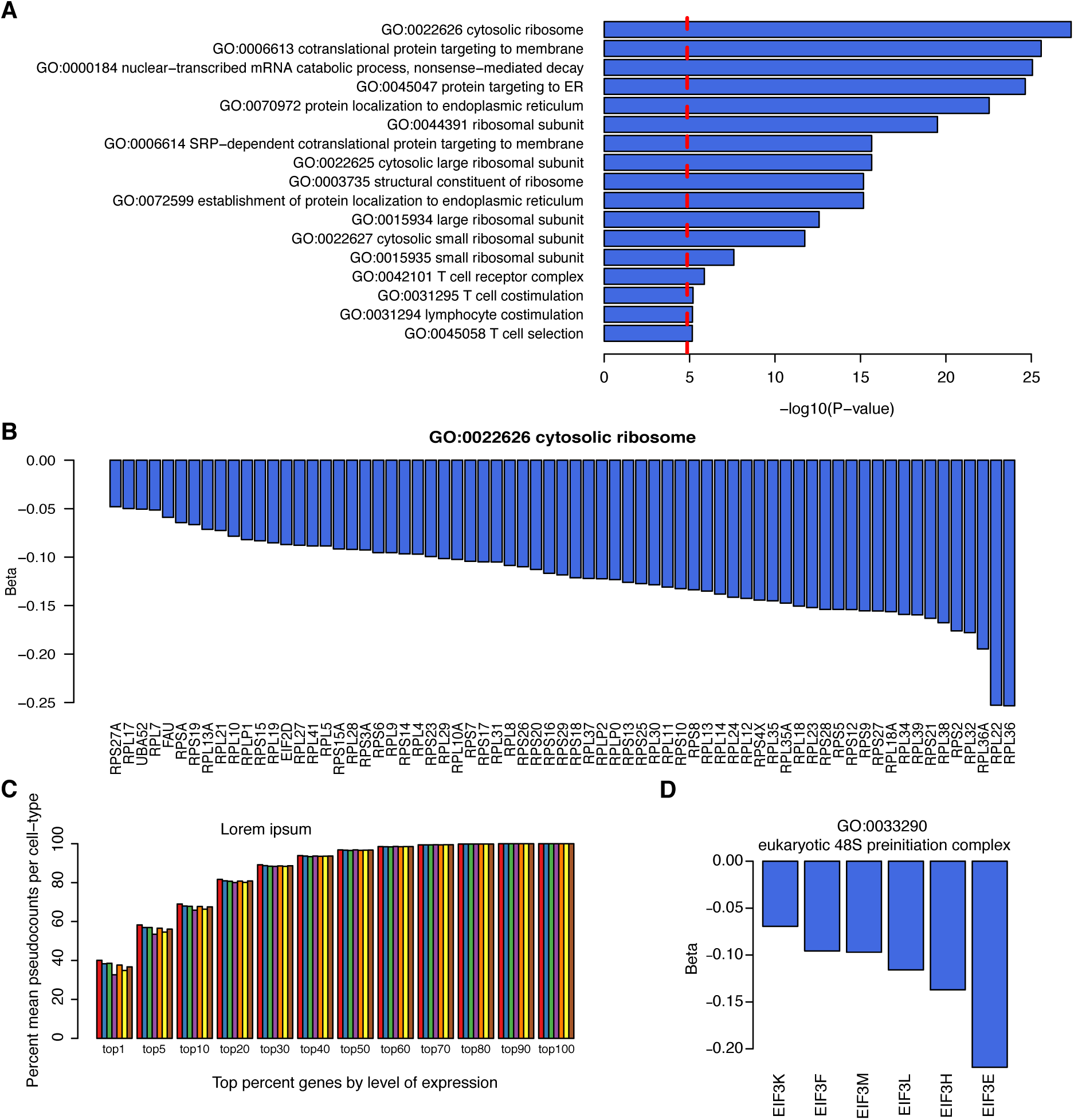
Ribosome associations with adaptiveness. (A) GO terms significantly enriched within adaptiveness-associated genes. (B) Innateness score (β) for genes with GO term cytosolic ribosome (GO:0022626) and with P < 2.5e-06. (C) Mean percent expression (Y-axis) occupied by the top X% expressed genes (X-axis) per cell-type. Cell types, from left to right are CD4+ T (red), CD8+ T (blue), MAIT (green), iNKT (purple), Vδ1 (orange), Vδ1 (yellow), NK (brown). (D) Innateness score (β) for genes with GO term eukaryotic 48S preinitiation complex (GO:0033290) and with P < 2.5e-06.

**Supplementary Figure 7.**
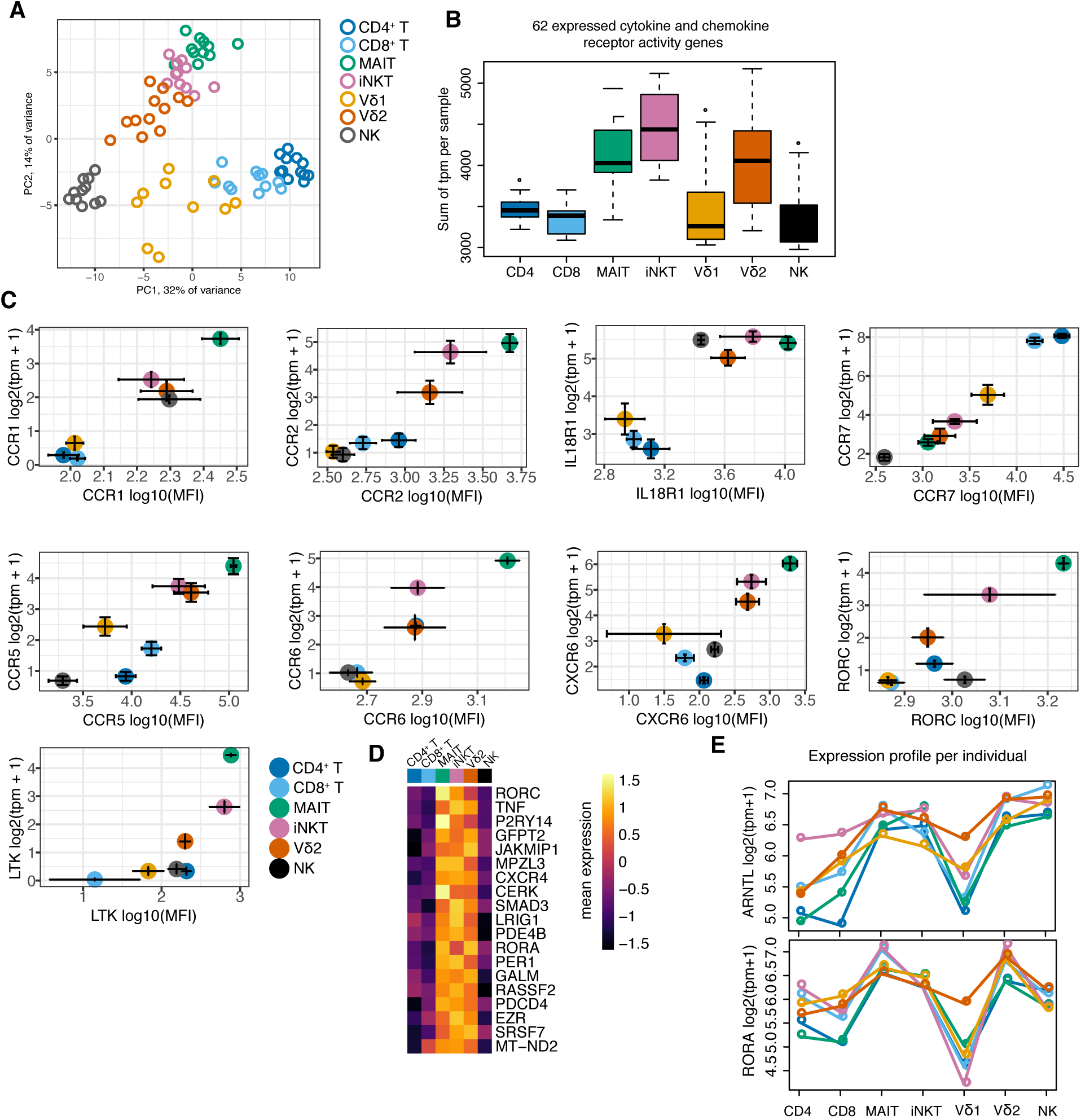
Transcription factors in ITCs. (A) Principal component analysis performed on 142 transcription factor genes variable among cell types (F statistic, P < 5.7e-05, Bonferroni threshold), centered and scaled genes. Plotted are scores for PC1 and PC2. (B) Sum of expression levels for 62 cytokine and chemokine receptor genes across samples.(C) Flow cytometric validation for genes differentially expressed between PLZF^+^ ITCs and adaptive T cells, showing protein levels by flow cytometry (X-axis), and transcript levels with RNA-seq (Y-axis). tpm, N=6; MFI, N=3. (D) Heatmap for mean expression level of PLZF target genes in mouse. Genes shown are upregulated in PLZF^+^ ITCs compared to adaptive T cells and low in NK cells. (E) Mean expression per individual (colored lines) among different cell types for circadian transcription factors ARNTL (that codes for BMAL, top) and RORA (bottom). All cell isolations were performed in the morning.

**Supplementary Figure 8.**
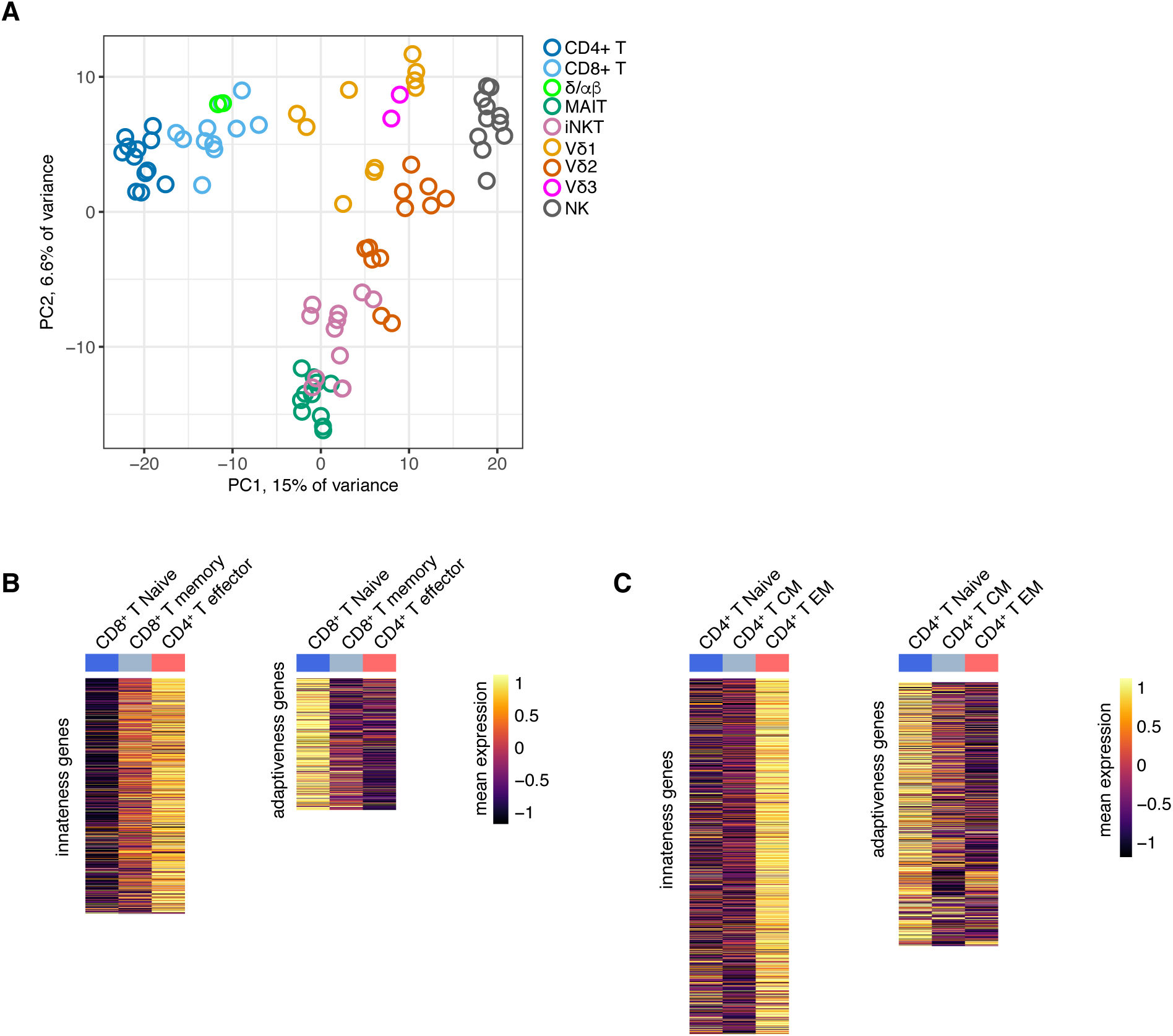
Innateness in candidate ITCs and adaptive T cells. (A) Principal component analysis including δα/β and Vδ3 T cells, performed on the top 1012 variable and expressed genes, centered and scaled. Plotted are scores for PC1, PC2. Heatplots of mean expression per cell-type for genes associated with innateness (left) or adaptiveness (right), for (B) HCMV specific CD8^+^ T cells and (C) CD4^+^ T cells.

**Supplementary Figure 9.**
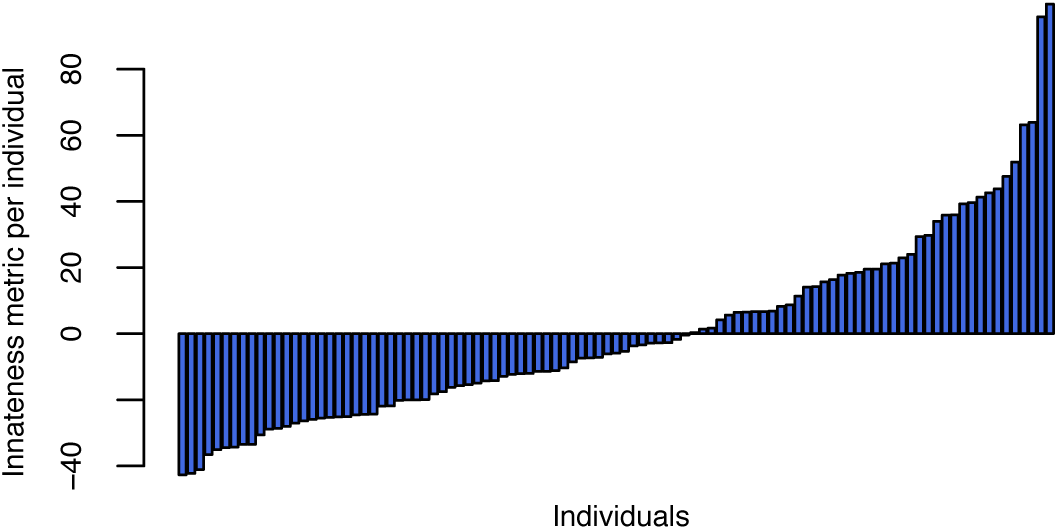
Individual innateness metric. Innateness metric calculated per individual by integrating the immunoprofiling data with the innateness gradient rank per cell type. Specifically, we summed the abundance per cell type (proportion of T cells) multiplied by the rank of that cell type in the innateness gradient. We then regressed out the age effects. Plotted are the residuals of this regression.

## References

1. P. J. Brennan, M. Brigl,M. B. Brenner, Invariant natural killer T cells: an innate activation scheme linked to diverse effector functions. Nat Rev Immunol 13, 101 (Feb, 2013).

2. D. I. Godfrey, A. P. Uldrich, J. McCluskey, J. Rossjohn, D. B. Moody, The burgeoning family of unconventional T cells. Nat Immunol 16, 1114 (Nov, 2015).

3. M. Salio, J. D. Silk, E. Y. Jones, V. Cerundolo, Biology of CD1- and MR1-restricted T cells. Annu Rev Immunol 32, 323 (2014).

4. I. Van Rhijn, D.B. Moody, Donor Unrestricted T Cells: A Shared Human T Cell Response. J Immunol 195, 1927 (Sep 1, 2015).

5. A. Bendelac, Positive selection of mouse NK1+ T cells by CD1-expressing cortical thymocytes. J Exp Med 182, 2091 (Dec 1, 1995).

6. A. Bendelac et al., CD1 recognition by mouse NK1+ T lymphocytes. Science 268, 863 (May 12, 1995).

7. M. Exley, J. Garcia, S. P. Balk, S. Porcelli, Requirements for CD1d recognition by human invariant Valpha24+ CD4-CD8-T cells. J Exp Med 186, 109 (Jul 7, 1997).

8. L. Kjer-Nielsen et al., MR1 presents microbial vitamin B metabolites to MAIT cells. Nature 491, 717 (Nov 291, 2012).

9. E. Treiner et al., Selection of evolutionarily conserved mucosal-associated invariant T cells by MR1. Nature 422, 164 (apr 13, 2003).

10. C. Harly et al., Key implication of CD277/butyrophilin-3 (BTN3A) in cellular stress sensing by a major human gammadelta T-cell subset. Blood 120, 2269 (Sep 13, 2012).

11. A. Palakodeti et al., The molecular basis for modulation of human Vgamma9Vdelta2 T cell responses by CD277/butyrophilin-3 (BTN3A)-specific antibodies. J Biol Chem 287, 32780 (Sep 21, 2012). 12.

12. S. Vavassori et al., Butyrophilin 3A1 binds phosphorylated antigens and stimulates human gammadelta T cells. Nat Immunol 14, 908 (Sep, 2013).

13. H. Wang et al., Butyrophilin 3A1 plays an essential role in prenyl pyrophosphate stimulation of human Vgamma2Vdelta2 T cells. J Immunol 191, 1029 (Aug 01, 2013).

14. A. Hayday, P. Vantourout, A long-playing CD about the gammadelta TCR repertoire. Immunity 39, 994 (Dec 12, 2013).

15. R. Di Marco Barros et al., Epithelia Use Butyrophilin-like Molecules to Shape Organ-Specific gammadelta T Cell Compartments. Cell 167, 203 (Sep 22, 2016).

16. A. C. Kohlgruber, C. A. Donado, N. M. LaMarche, M. B. Brenner, P. J. Brennan, Activation strategies for invariant natural killer T cells. Immunogenetics, (Jul 25, 2016).

17. M. Brigl, L. Bry, S. C. Kent, J. E. Gumperz, M.B. Brenner, Mechanism of CD1d-restricted natural killer T cell activation during microbial infection. Nat Immunol 4, 1230 (Dec, 2003).

18. A. J. Tyznik et al., Cutting edge: the mechanism of invariant NKT cell responses to viral danger signals. J Immunol 181, 4452 (Oct 1, 2008).

19. J. E. Ussher et al., CD161++ CD8+ T cells, including the MAIT cell subset, are specifically activated by IL-12+IL-18 in a TCR-independent manner. Eur J Immunol 44, 195 (Jan, 2014).

20. B. van Wilgenburg et al., MAIT cells are activated during human viral infections. Nat Commun 7, 11653 (Jun 23, 2016). 29

21. A. Cesano, S. Visonneau, S. C. Clark, D. Santoli, Cellular and molecular mechanisms of activation of MHC nonrestricted cytotoxic cells by IL-12. J Immunol 151, 2943 (Sep 15, 1993).

22. H. DasM. Sugita, M. B. Brenner, Mechanisms of Vdelta1 gammadelta T cell activation by microbial components. J Immunol 172, 6578 (Jun, 2004).

23. J. L. Klein, H. Fickenscher, J. E. Holliday, B. Biesinger, B. Fleckenstein, Herpesvirus saimiri immortalized gamma delta T cell line activated by IL-12. J Immunol 156, 2754 (Apr 15, 1996).

24. C. Ueta et al., Interleukin-12 activates human gamma delta T cells: synergistic effect of tumor necrosis factor-alpha. Eur J Immunol 26, 3066 (Dec, 1996).

25. J. Novak, J. Dobrovolny, L. Novakova, T. Kozak, The decrease in number and change in phenotype of mucosal-associated invariant T cells in the elderly and differences in men and women of reproductive age. Scandinavian journal of immunology 80, 271 (Oct, 2014).

26. E. Patin et al., Natural variation in the parameters of innate immune cells is preferentially driven by genetic factors. Nat Immunol 19, 302 (Mar, 2018).

27. L. A. O’Neill, R. J. Kishton, J. Rathmell,A guide to immunometabolism for immunologists. Nat Rev Immunol 16, 553 (Sep, 2016).

28. J. van Riggelen, A. Yetil, D. W. Felsher, MYC as a regulator of ribosome biogenesis and protein synthesis. Nature reviews. Cancer 10, 301 (Apr, 2010).

29. A. David et al., RNA binding targets aminoacyl-tRNA synthetases to translating ribosomes. J Biol Chem 286, 20688 (Jun 10, 2011). 30

30. A. N. Milne, W. W. Mak, J. T. Wong, Variation of ribosomal proteins with bacterial growth rate. J Bacteriol 122, 89 (Apr, 1975).

31. E. Bosdriesz, D. Molenaar, B. Teusink, F. J. Bruggeman, How fast-growing bacteria robustly tune their ribosome concentration to approximate growth-rate maximization. The FEBS journal 282, 2029 (May, 2015).

32. H. Y. Kim et al., The development of airway hyperreactivity in T-bet-deficient mice requires CD1d-restricted NKT cells. J Immunol 182, 3252 (Mar 1, 2009).

33. S. J. Szabo et al., A novel transcription factor, T-bet, directs Th1 lineage commitment. Cell 100, 655 (Mar 17, 2000).

34. S. J. Szabo et al., Distinct effects of T-bet in TH1 lineage commitment and IFN-gamma production in CD4 and CD8 T cells. Science 295, 338 (Jan 11, 2000.

35. I. Albrecht et al., Persistence of effector memory Th1 cells is regulated by Hopx. Eur J Immunol 40, 2993 (Nov, 2010). 35.

36. C. X. Dominguez et al., The transcription factors ZEB2 and T-bet cooperate to program cytotoxic T cell terminal differentiation in response to LCMV viral infection. J Exp Med 212, 2041 (Nov 16, 2000).

37. K. D. Omilusik et al., Transcriptional repressor ZEB2 promotes terminal differentiation of CD8+ effector and memory T cell populations during infection. J Exp Med 212, 2027 (Nov 16, 2000).

38. B. N. Weber et al., A critical role for TCF-1 in T-lineage specification and differentiation. Nature 476, 63 (Aug 3, 2011).

39. L. M. D’Cruz, M. H. Stradner, C. Y. Yang, A. W. Goldrath, E and Id proteins influence invariant NKT cell sublineage differentiation and proliferation. J Immunol 192, 2227 (Mar 1, 2014).

40. M. Kanda et al., Transcriptional regulator Bhlhe40 works as a cofactor of T-bet in the regulation of IFN-gamma production in iNKT cells. Proc Natl Acad Sci U S A 113, E3394 (Jun 14, 2016).

41. V. J. Lawson, D. Maurice, J. D. Silk, V. Cerundolo, K. Weston, Aberrant selection and function of invariant NKT cells in the absence of AP-1 transcription factor Fra-2. J Immunol 183, 2575 (Aug 15, 2009).

42. L. A. Monticelli et al., Transcriptional regulator Id2 controls survival of hepatic NKT cells. Proc Natl Acad Sci U S A 106, 19461 (Nov 17, 2009).

43. M. Verykokakis et al., Essential functions for ID proteins at multiple checkpoints in invariant NKT cell development. J Immunol 191, 5973 (Dec 15, 2013).

44. H. Spits, J. P. Di Santo, The expanding family of innate lymphoid cells: regulators and effectors of immunity and tissue remodeling. Nat Immunol 12, 21 (Jan, 2011).

45. M. H. Stradner, K. P. Cheung, A. Lasorella, A. W. Goldrath, L. M. D’Cruz, Id2 regulates hyporesponsive invariant natural killer T cells. Immunol Cell Biol 94, 640 (Aug, 2016).

46. W. K. Mowel et al., Group 1 Innate Lymphoid Cell Lineage Identity Is Determined by a cis-Regulatory Element Marked by a Long Non-coding RNA. Immunity 47, 435 (Sep 19, 2017).

47. F. Masson et al., Id2-mediated inhibition of E2A represses memory CD8+ T cell differentiation. J Immunol 190, 4585 (May 1, 2013). 3

48. R. B. Delconte et al., The Helix-Loop-Helix Protein ID2 Governs NK Cell Fate by Tuning Their Sensitivity to Interleukin-15. Immunity 44, 103 (Jan 19, 2016).

49. D. Kovalovsky et al., The BTB-zinc finger transcriptional regulator PLZF controls the development of invariant natural killer T cell effector functions. Nat Immunol 9, 1055 (Sep, 2008).

50. A. K. Savage et al., The transcription factor PLZF directs the effector program of the NKT cell lineage. Immunity 29, 391 (Sep 19, 2008).

51. M. G. Constantinides, B. D. McDonald, P. A. Verhoef, A. Bendelac, A committed precursor to innate lymphoid cells. Nature 508, 397 (Apr 17, 2014).

52. M. Eidson et al., Altered development of NKT cells, gammadelta T cells, CD8 T cells and NK cells in a PLZF deficient patient. PLoS One 6, e24441 (2011).

53. A. P. Mao et al., Multiple layers of transcriptional regulation by PLZF in NKT-cell development. Proc Natl Acad Sci U S A 113, 7602 (Jul 05, 2016).

54. G. E. Duffield et al., A role for Id2 in regulating photic entrainment of the mammalian circadian system. Curr Biol 19, 297 (Feb 24, 2009).

55. S. Honma et al., Dec1 and Dec2 are regulators of the mammalian molecular clock. Nature 419, 841 (Oct 24, 2002).

56. S. M. Ward, S. J. Fernando, T. Y. Hou, G.E. Duffield, The transcriptional repressor ID2 can interact with the canonical clock components CLOCK and BMAL1 and mediate inhibitory effects on mPer1 expression. J Biol Chem 285, 38987 (Dec 10, 2010).

57. B. A. Mangan et al.,Cutting edge: CD1d restriction and Th1/Th2/Th17 cytokine secretion by human Vdelta3 T cells. J Immunol 191, 30 (Jul 1, 2013). 33

58. D. G. Pellicci et al., The molecular bases of delta/alphabeta T cell-mediated antigen recognition. J Exp Med 211, 2599 (Dec 15, 2014).

59. T. Kawabe et al., Memory-phenotype CD4(+) T cells spontaneously generated under steady-state conditions exert innate TH1-like effector function. Science immunology 2, (Jun 16, 2017).

60. K. M. Hertoghs et al., Molecular profiling of cytomegalovirus-induced human CD8+ T cell differentiation. J Clin Invest 120, 4077 (Nov, 2010).

61. V. Ranzani et al., The long intergenic noncoding RNA landscape of human lymphocytes highlights the regulation of T cell differentiation by linc-MAF-4. Nat Immunol 16, 318 (Mar, 2015).

62. L. Guo et al., Innate immunological function of TH2 cells in vivo. Nat Immunol 16, 1051 (Oct, 2015).

63. A. Mitson-Salazar et al., Hematopoietic prostaglandin D synthase defines a proeosinophilic pathogenic effector human T(H)2 cell subpopulation with enhanced function. J Allergy Clin Immunol 137, 907 (Mar, 2016).

64. B. Upadhyaya, Y. Yin, B. J. Hill, D. C. Douek, C. Prussin, Hierarchical IL-5 expression defines a subpopulation of highly differentiated human Th2 cells. J Immunol 187, 3111 (Sep 15, 2011).

65. E. Wambre et al., A phenotypically and functionally distinct human TH2 cell subpopulation is associated with allergic disorders. Science translational medicine 9, (Aug 2, 2017). 34

66. M. S. Davey et al., Clonal selection in the human Vdelta1 T cell repertoire indicates gammadelta TCR-dependent adaptive immune surveillance. Nat Commun 8, 14760 (Mar 1, 2017).

67. Y. Liu, J. Zhou, K.P. White, RNA-seq differential expression studies: more sequence or more replication? Bioinformatics 30, 301 (Feb 1, 2014).

68. S. Picelli et al., Full-length RNA-seq from single cells using Smart-seq2. Nature protocols 9, 171 (Jan, 2014).

69. N. L. Bray, H. Pimentel, P. Melsted, L. Pachter, Near-optimal probabilistic RNA-seq quantification. Nat Biotechnol 34, 525 (May, 2016).

70. Expansion of the Gene Ontology knowledgebase and resources. Nucleic Acids Res 45, D331 (Jan 4, 2017).

71. M. Ashburner et al., Gene ontology: tool for the unification of biology. The Gene Ontology Consortium. Nat Genet 25, 25 (May, 2000).

72. E. Eden, D. Lipson, S. Yogev, Z. Yakhini, Discovering motifs in ranked lists of DNA sequences. PLoS computational biology 3, e39 (Mar 23, 2007).

73. M. Kanehisa, S. Goto, KEGG: kyoto encyclopedia of genes and genomes. Nucleic Acids Res 28, 27 (Jan 1, 2000).

74. A. Kamburov, C. Wierling, H. Lehrach, R. Herwig, ConsensusPathDB--a database for integrating human functional interaction networks. Nucleic Acids Res 37, D623 (Jan, 2009).

75. D. Smedley et al., The BioMart community portal: an innovative alternative to large, centralized data repositories. Nucleic Acids Res 43, W589 (Jul 1, 2015). 35

76. N. R. Cohen et al., Shared and distinct transcriptional programs underlie the hybrid nature of iNKT cells. Nat Immunol 14, 90 (Jan, 2013).

77. T. S. Heng, M. W. Painter, The Immunological Genome Project: networks of gene expression in immune cells. Nat Immunol 9, 1091 (Oct, 2008).

